# Reticular adhesions: A new class of adhesion complex that mediates cell-matrix attachment during mitosis

**DOI:** 10.1101/234237

**Authors:** John G. Lock, Matthew C. Jones, Janet A. Askari, Xiaowei Gong, Anna Oddone, Helene Olofsson, Sara Göransson, Melike Lakadamyali, Martin J. Humphries, Staffan Strömblad

**Author notes:** These authors contributed equally to this study (Matthew C. Jones, Janet A. Askari, Xiaowei Gong). Correspondence: Staffan Strömblad, Karolinska Institutet, Novum, Hälsovägen 7-9, SE-141 83 Huddinge, Sweden. John Lock, University of New South Wales, School of Medical Sciences, Kensington 2052 Sydney, Australia.

## Abstract

Adhesion to the extracellular matrix (ECM) persists during mitosis in most cell types. Yet, classical adhesion complexes (ACs), such as focal adhesions and focal complexes, do and must disassemble to enable cytoskeletal rearrangements associated with mitotic rounding. Given this paradox, mechanisms of mitotic cell-ECM adhesion remain undefined. Here, we identify ‘reticular adhesions’, a new class of AC that is mediated by integrin αvβ5, formed during interphase and preserved at cell-ECM attachment sites throughout cell division. Consistent with this role, integrin β5 depletion perturbs mitosis and disrupts spatial memory transmission between cell generations. Quantitative imaging reveals reticular adhesions to be both morphologically and dynamically distinct from classic focal adhesions, while mass spectrometry defines their unique composition; lacking virtually all consensus adhesome components. Indeed, remarkably, reticular adhesions are functionally independent of both talin and F-actin, yet are promoted by phosphatidylinositol-4,5-bisphosphate (PI-4,5-P2). Overall, the distinct characteristics of reticular adhesions provide a unique solution to the problem of maintaining cell-ECM attachment during mitotic rounding and division.

## Introduction

Cell-to-ECM attachment is primarily achieved through a range of integrin-containing ACs, including focal complexes, focal adhesions and fibrillar adhesions (Balaban et al., 2001; Hynes, 2002; Lock et al., 2008). This attachment modulates many processes, including cell movement, differentiation and proliferation (Berrier and Yamada, 2007). Though structurally and functionally varied, ACs overlap substantially in their molecular composition, sharing a consensus adhesome of approximately 60 proteins (Horton et al., 2015). As one of the most abundant consensus adhesome proteins, talin is commonly viewed as an indispensable contributor to integrin activation (Tadokoro et al., 2008), AC development (Changede et al., 2015) and AC organisation (Liu et al., 2015). Adaptor proteins that couple integrins to F-actin, such as vinculin and paxillin, are also universally associated with ACs, reflecting the pivotal role of F-actin in AC maturation, maintenance and function (Gardel et al., 2010).

Although ACs have mostly been studied during interphase, adhesion is necessary for mitotic progression and for the transmission of spatial memory between cell generations (Jime et al., 2007; Minc et al., 2011), a key factor controlling differentiation and tissue development (Akanuma et al., 2016). Paradoxically, the importance of cell-ECM adhesion during mitosis conflicts with the observed disassembly of ACs during mitotic onset (Maddox and Burridge, 2003), and with evidence that failure of AC disassembly perturbs cell division (Dao et al., 2009; Lancaster et al., 2013). Furthermore, integrins implicated in mitotic adhesion, such as β1, appear to function not at the adhesion plane, but in the detached cell cortex (Petridou and Skourides, 2016). Overall, the nature of mitotic ACs remains profoundly unclear (LaFlamme et al., 2008; Ramkumar and Baum, 2016).

Here, we describe a new class of ‘reticular’ AC with a unique adhesome, formed by integrin αVβ5 during interphase in the absence of both talin and F-actin. Reticular ACs persist throughout mitosis, providing ECM anchoring that is necessary for efficient cell division. Thus, reticular adhesions not only constitute a novel adhesion type, but provide a unique solution to the paradox of mitotic cell-ECM attachment.

## Results

### αVβ5 is the predominant integrin subunit within ACs in cells grown in long-term culture

The well-defined integrin consensus adhesome is derived from cells plated for several hours on fibronectin (Horton et al., 2015; Winograd-Katz et al., 2014). To study the adhesome of proliferating cells in longer-term culture, we performed mass spectrometry analysis of integrin-associated AC composition in U2OS osteosarcoma cells following 72 h growth. Unexpectedly, the most abundant integrin subunits identified were αV and β5, with much lower levels of β1, β3, β8, α5 and α3 (Fig.1A). Immunofluorescence analysis confirmed that very distinct αVβ5-positive ACs were visible in a range of cells in long-term culture, with little αVβ3 or β1 labelling of ACs detected in U2OS, A549 and A375 cells (Fig.1B and Supplementary Fig.1A). Notably, in all three cell lines, αVβ5 was present not only in classical focal adhesions at the cell periphery, but also in reticular structures across the cell body.

**Figure 1.**
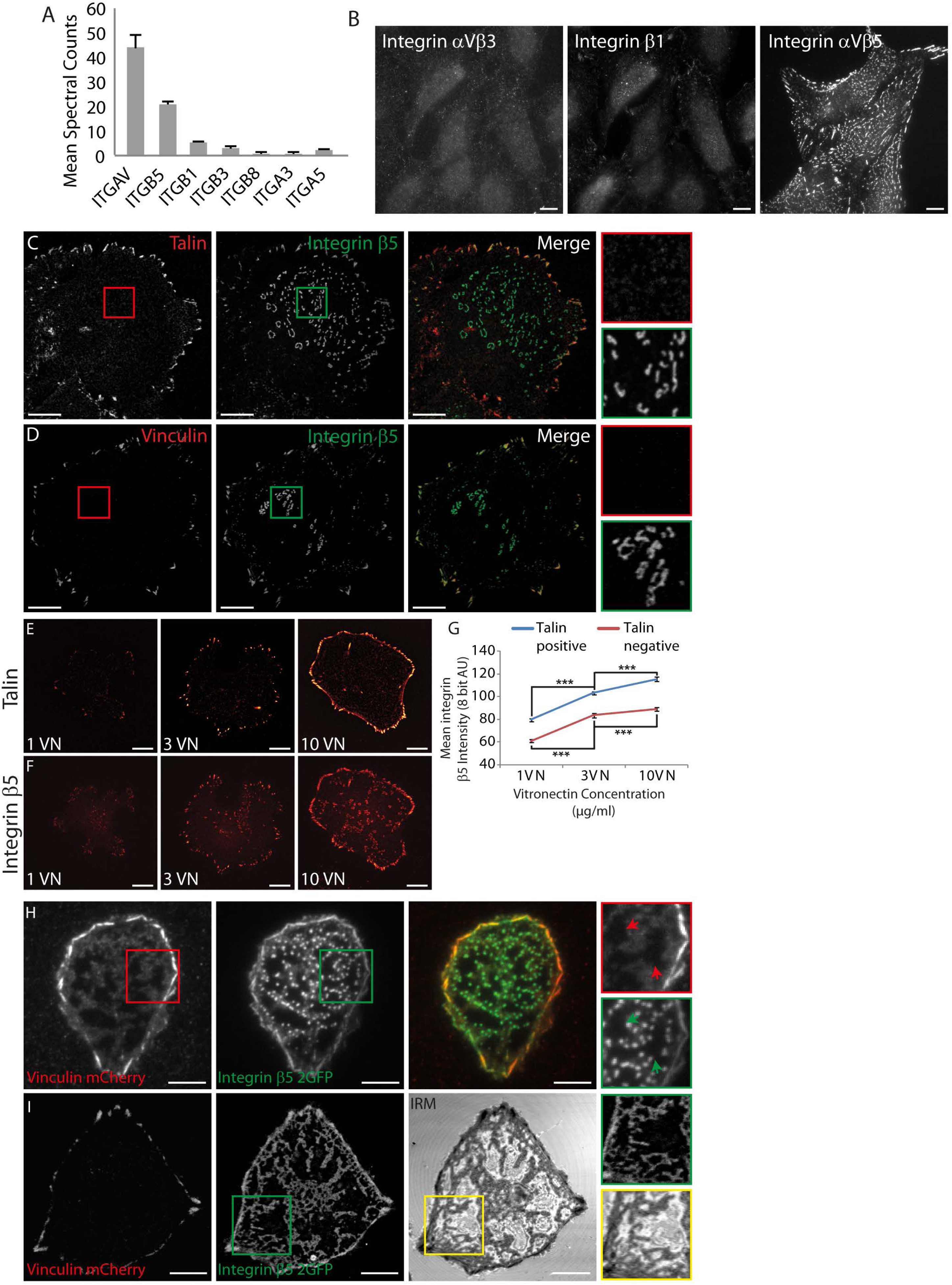
Integrin αVβ5 forms novel talin and vinculin negative reticular adhesion structures. (A) Mass spec analysis of integrin subunits detected in adhesions isolated (after cell removal) from U2OS cells plated in complete medium for 3 d on tissue culture plastic. Results are mean spectral counts of three separate experiments ± S.D. (B) U2OS cells were plated on glass coverslips for 72 h. Confocal images of immuno-fluorescently labeled integrins αVβ3 (LM609), β1 (9EG7) and αVβ5 (15F11). (C-F) U2OS cells plated for 3 h in serum-free media on surfaces coated with 10 μg/ml vitronectin (VN), except where otherwise specified. Confocal images of talin (C) or vinculin (D) immunofluorescence with that of β5. Boxed areas shown at higher magnification to the right (E,F) Co-labeling of talin (E) and β5 (F) in cells grown on glass coated with 1, 3, or 10 μg/ml VN. (G) Quantified intensities of talin-positive (blue) or –negative (red) β5 structures. Data from 81 cells (>23 per condition), 6132 adhesions. Error bars indicate 95% confidence intervals. Significant differences indicated (Mann-Whitney U test, *** = p < 0.001). (H) TIRF images of a vinculin mCherry and β5-2GFP-expressing U2OS cell (U2OS-β5 V). Arrows in magnified boxes highlight regions lacking vinculin signal, which fall between β5 -positive, vinculin-negative puncta. (I) Confocal and interference reflection microscopy (IRM) images of a U2OS-β5V cell exemplify correlations between β5-positive, vinculin-negative structures and regions with close cell-substrate proximity.

To characterise these reticular structures further, U2OS cells were plated on the integrin αV ligand vitronectin (VN). Confocal imaging of ventral membranes showed β5 associated with two different structures: peripheral ACs containing talin and vinculin, and centrally-distributed, punctate or reticular structures lacking these components (Fig.1C-D). Similar αVβ5-positive, talin-negative structures were detected in CS1-b5, HeLa, MCF7, and MAE cells (Supplementary Fig.1B). Integrin αV and β5 subunit colocalisation confirmed that these reticular structures contained both αVβ5 subunits (Supplementary Fig.2A). Co-labelling of β5 with antibodies directed against various AC-related proteins failed to reveal specific colocalisation with the reticular, β5-positive structures. Markers tested included consensus adhesome components, αVβ5-binding partners, cytoskeletal proteins including F-actin, and phosphotyrosine (Supplementary Fig.2B-M). Equivalent structures lacking F-actin are also shown in BT549 cells (Supplementary Fig.1B). Integrin β5 fluorescence intensity in both talin-positive and talin-negative structures correlated with VN concentrations (Fig.1E-G), while U2OS cells plated on laminin (not an αVβ5-ligand) only formed vinculin-positive ACs (Supplementary Fig.2N). These data demonstrate that formation of the reticular structures depends on αVβ5-ECM ligand binding.

We next expressed EGFP-tagged integrin β5 (integrin β5-2GFP) in U2OS cells and antibody labeled the integrin β5 extracellular domain without prior cell permeabilisation (Supplementary Fig.2O). Strong GFP-to-antibody colocalisation demonstrated αVβ5 plasma membrane embedding and antibody specificity. Moreover, total internal reflection (TIRF) imaging of live U2OS cells coexpressing β5-2GFP and vinculin-mCherry (U2OS-B5V) revealed central, αVβ5-positive, vinculin-negative structures in the TIRF plane (Fig.1H). Dark intracellular regions in vinculin-mCherry signals indicated where tensioned ventral membranes arced out of the TIRF plane, leaving no cytoplasmic signal. These dark regions corresponded with large gaps between αVβ5-postive, vinculin-negative puncta, suggesting them to be attachment points that pin the ventral plasma membrane to the substrate. This hypothesis was supported by live cell interference reflection microscopy, where close cell-substrate proximity corresponded precisely with integrin β5-2GFP signals in both vinculin-positive focal adhesions and vinculin-negative structures (Fig. 1I). Collectively, these data indicate that αVβ5-positive, consensus adhesome component-negative reticular structures are *bonafide* cell-ECM ACs. These are hereafter termed “reticular ACs”.

### Reticular and focal ACs are morphologically and dynamically distinct

Reticular ACs were more numerous than classical focal ACs at all sizes (Fig.2A), increased in size more frequently (Fig.2B) and were generally localised further from the cell periphery (Fig.2C). There was no correlation between reticular AC size and integrin β5 clustering density, unlike the increased integrin density observed in larger focal ACs (Fig.2D) (Hernández-Varas et al., 2015; Kiss et al., 2015; Lock et al., 2014). This implies molecular-scale differences between the maturation of focal and reticular adhesions, with the latter being more homogenous. Reticular adhesions formed as small puncta, grew by net peripheral integrin recruitment, producing ring-like or reticular structures that ultimately fragmented and disassembled, all without recruiting vinculin (Fig.2E-H; Supplementary Movie 1 and cropped region from Fig.2H in Supplementary Movie 2). Thus, reticular adhesions form *de novo* as a distinct class of AC independently from focal adhesions.

**Figure 2.**
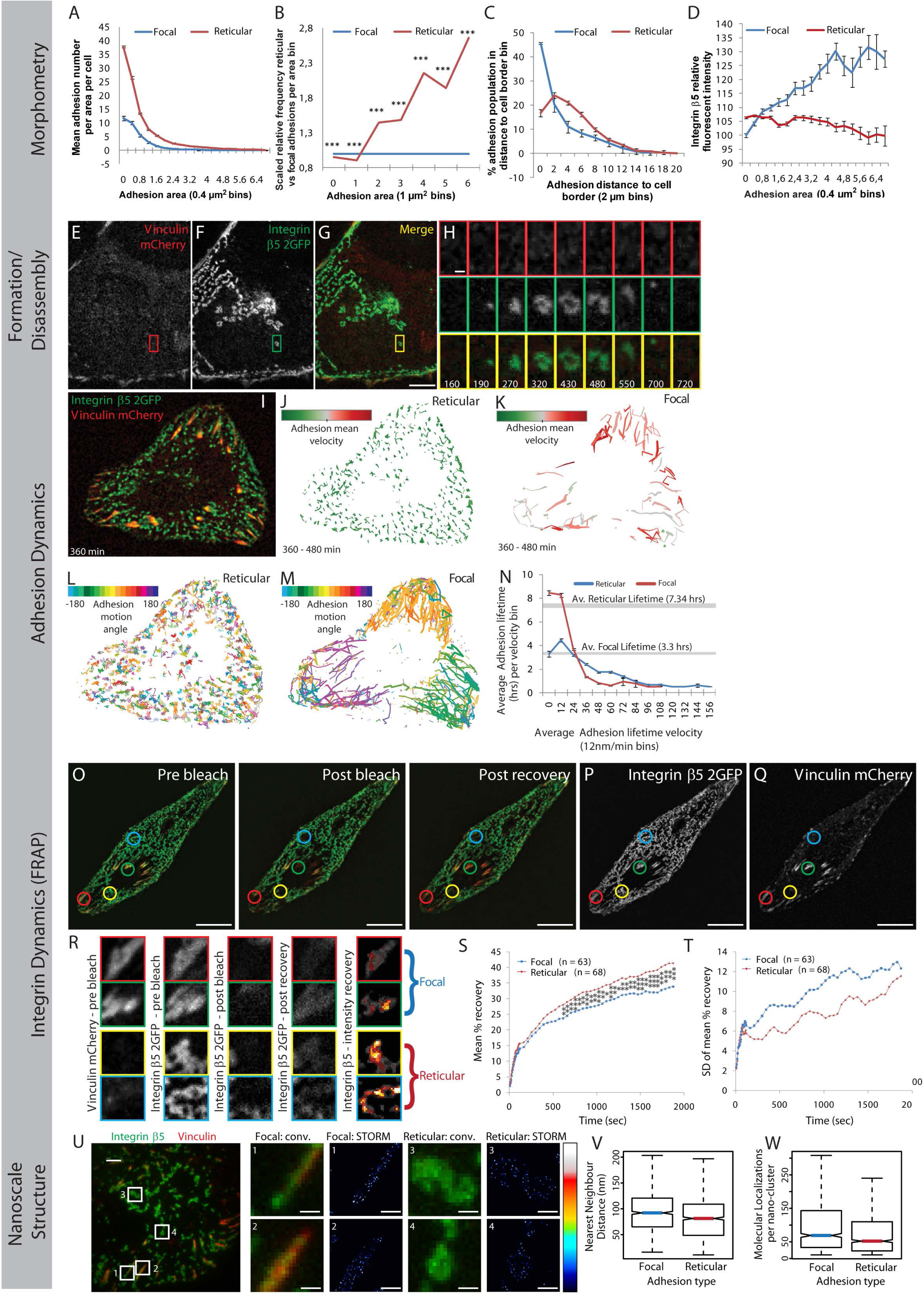
Comparison of focal and reticular adhesion morphometry and dynamics. (A) Histogram of focal (blue) and reticular adhesions (red) by area, per U2OS cell (error bars = 95% confidence intervals (CI)). (B) Frequency of reticular adhesions by area, scaled by total of each adhesion class, represented as fold-change relative to focal adhesions. Significant differences indicated (Mann-Whitney U test, *** = p < 0.001). (C) Percentage of adhesions of each class located at various distances from cell border (error bars = 95% CI). (D) Adhesion area *versus* mean integrin β5 intensity, scaled relative to smallest detectable focal adhesions (error bars = 95% CI). (E) Representative image of Vinculin mCherry and (F) β5-2GFP (merged in G; see **Supplementary Movie 1**). (H) Zoomed regions cropped from E-G at time points indicated (160-720 min relative to imaging start; see **Supplementary Movie 2**). Scale bars: E-G (10 μm); H (1 μm). (I-M, see **Supplementary Movie 3**) (I) Merged image of β5-2GFP and vinculin mCherry at representative timepoint (360 min). Trajectories of reticular (J) and focal adhesions (K) colour-coded by mean velocity (green = slow; red = fast). Trajectories of reticular (L) and focal adhesions (M) colour-coded (as indicated) by net adhesion motion angle. Line thicknesses in J-M indicate instantaneous adhesion velocity. (N) Aggregate analysis of all trajectories of average adhesion velocity (over lifetime) *versus* corresponding average adhesion lifetime (hours; dashed lines indicate adhesion class average lifetimes; error bars indicate = 95% CI). Data in A-D and N derive from live imaging and analysis of 14 U2OS-β5V cells over 12 h (10 min intervals), providing 30 123 focal adhesion and 91 898 reticular adhesion observations from the individual lifetimes of >1500 focal and >2000 reticular adhesions. (O-Q, see **Supplementary Movie 4**) (O) Integrin β5-2GFP (green) and vinculin mCherry (red) in a representative cell pre-bleach, immediately post-bleach and post recovery (30 min). Circles indicate bleach regions: green and red circles target focal adhesions; blue and yellow circles target reticular adhesions, is clearly shown in single channel images (P-Q). (R) Square regions corresponding to circles in P-Q show vinculin mCherry adhesion signals pre-bleach (1st column), and β52GFP signals pre-bleach (2^nd^ column), post-bleach (3^rd^ column) and post-recovery (4^th^ column). Colour-scaled images (low to high values = black, red, orange, yellow, white) calculated by subtracting post bleach intensities from post recovery intensities for focal (upper two rows) and reticular (lower two rows) adhesions (5th column). (S) Aggregate FRAP recovery curves (postbleach recovery time *versus* mean fluorescence recovery as percentage of pre-bleach intensity) for 63 focal and 68 reticular adhesions (from 15 cells). Differences in recovery extent and rate apparent from ~600 s based on Mann-Whitney U test, *** = p < 0.001, ** = p < 0.01. (T) Post-bleach recovery time versus standard deviation of recovery at each time point. Scale bars: O-Q (10 μm). (UW) Comparison of integrin β5 nanoclustering. (U) Integrin β5 immuno-labeling and vinculin mCherry in a representative U2OS cell plated for 3 h on VN and imaged via conventional (diffraction-limited) confocal microscopy. Representative focal (1 and 2) and reticular adhesions (3 and 4) cropped from matched conventional (conv., left, vinculin and β5) and stochastic optical reconstruction microscopy (STORM, β5 only, ‘royal’ look-up table scales from minimum (black) to maximum (white) intensity values as indicated in legend) images (right). Scale bars 2 μm (500 nm in cropped images). Quantification of integrin β5 nanocluster nearest neighbour distances (V) and molecular localization counts per nanocluster (W) based on STORM data. In total, 216 focal and 162 reticular adhesions were assessed, including > 5500 nanoclusters. Boxplot notches approximate 95% CI (see methods for all details).

Quantitative adhesion tracking highlighted stark differences in dynamics between reticular and focal AC structures (Fig.2I-N; Supplementary Movie 3): isotropic reticular AC growth produced low displacement (Fig.2J), while focal ACs elongated anisotropically and slid at high velocities, reflecting F-actin-derived forces driving asymmetric component recruitment (Fig.2K) (Ballestrem et al., 2001; Besser and Safran, 2006). Isotropic growth and immobility in reticular ACs suggests the absence of such directed mechanical cues (Yu et al., 2013) and complements the observed lack of F-actin at these ACs. This conclusion was further supported by the locally disordered motion of reticular AC trajectories (Fig.2L). In contrast, focal ACs moved co-linearly within different cell lobes (Fig.2M), reflecting aligned, centripetal F-actin-derived forces (Mohl et al., 2012). The relationship between average AC velocity and AC lifetime revealed that for both focal and reticular ACs, fast movement corresponded with short lifetime. Thus, fast-moving focal ACs existed for less than half the lifespan of reticular ACs, which were relatively static and long-lived (Fig.2N).

Fluorescence recovery after photobleaching (FRAP) analysis revealed that despite their increased lifetime as complexes, β5-2GFP turnover in reticular ACs was both faster and more extensive than in focal ACs (Fig.2O-T; Supplementary Movie 4). Conversely, variability in β5-2GFP fluorescence recovery was lower in reticular ACs (Fig.2T), suggesting relative homogeneity in their molecular organisation and dynamics, including across their lifespan. This corresponds with the homogeneity in integrin clustering densities (lack of dependence on AC size) also observed in reticular ACs (Fig.2D).

Stochastic optical reconstruction microscopy (STORM) showed that both AC types consist of small internal clusters, here termed ‘nanoclusters’ (Fig.2U). Minimal differences were observed between the AC types in terms of nearest neighbour distances between integrin β5 nanoclusters, or molecular localisation counts per nanocluster (Fig.2V-W). Thus, despite the absence of consensus adhesome components like talin – thought to control nanoscale integrin organisation (Liu et al., 2015) – and differences in macromolecular dynamics, the molecular scale organisation of integrin β5 is virtually identical in focal and reticular ACs.

### Reticular ACs mediate cell attachment but are functionally independent of F-actin and talin

Although F-actin and talin were not detected in reticular ACs, we tested whether these components may be functionally significant at levels below detection limits (Fig.3; Supplementary Fig.3). Disruption of actin polymerisation by cytochalasin D caused disassembly of focal ACs but retention of reticular ACs. Cytochalasin D treatment prior to cell-ECM attachment inhibited focal adhesion formation, while reticular ACs still formed (Fig.3A-B; Supplementary Fig.3A-B and D-E and Supplementary Movie 5). Inhibition of actin polymerisation using latrunculin A produced equivalent results (Supplementary Fig.3C). Cytochalasin D inhibited cell spreading, but not reticular AC numbers relative to cell area, as evidenced by matched linear trends in cell area *versus* AC number within treated and control cells (Fig.3C). Notably, while cytochalasin D substantially reduced vinculin levels in surviving focal ACs, β5 densities actually increased in reticular ACs (Fig.3D).

**Figure 3.**
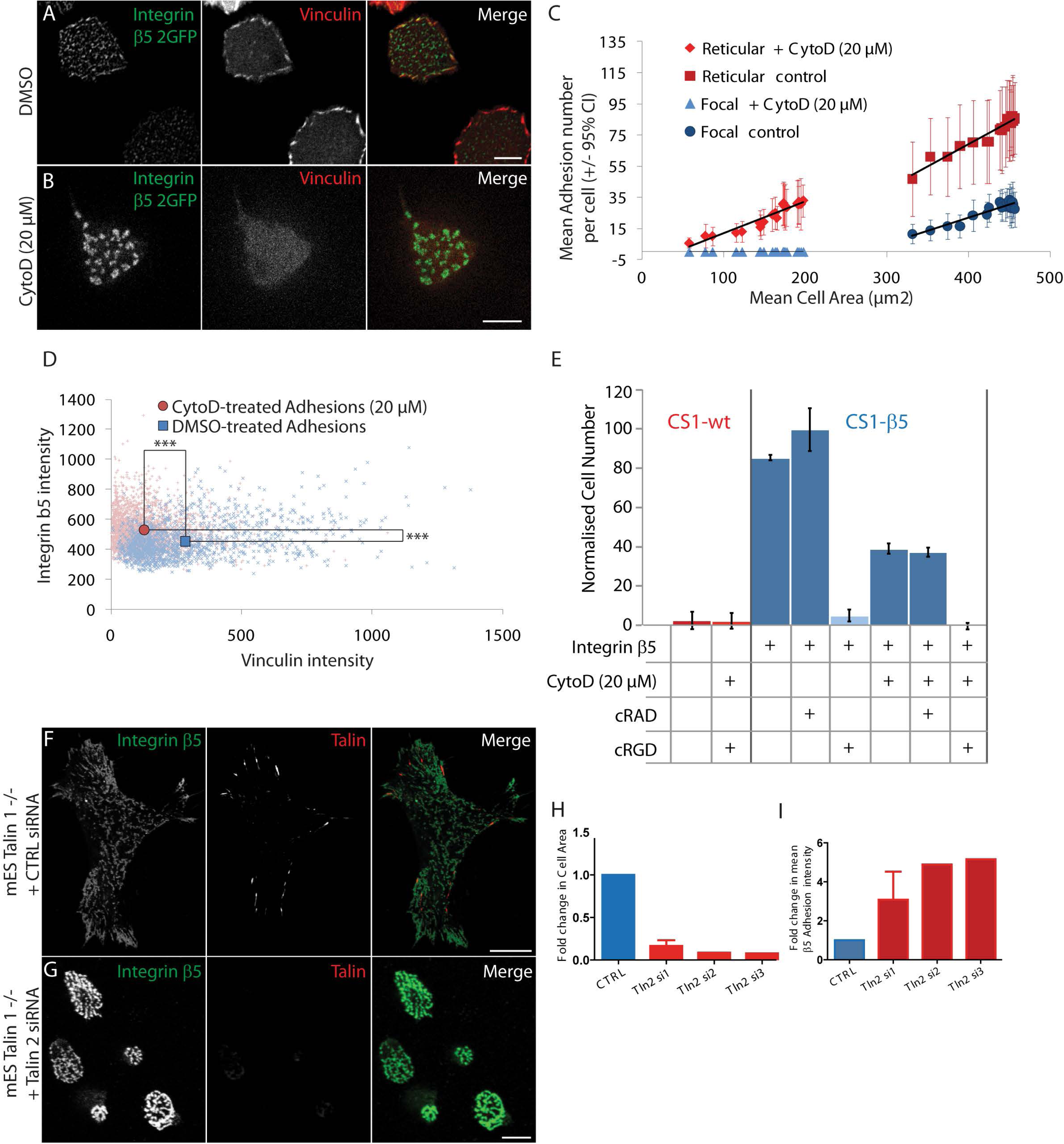
Reticular adhesions form in the absence of F-actin and talin. (A-B) Confocal images of integrin β5-2GFP and vinculin mCherry in live U2OS β5 V cells 1.8 h post attachment to vitronectin (VN)-coated surfaces (see **Supplementary Movie 5**). Cells pre-treated in suspension (30 min) and during spreading with DMSO (A) or 20 μM of F-actin polymerization inhibitor cytochalasin D (CytoD) (B). (C) Comparison of mean cell area μm^2^, per cell over 4 h time course) *versus* mean reticular (red) or focal (blue) adhesion number (per cell over 4 h time course) following treatments (black lines = linear regression, error bars = 95% CI, 12 cells per condition, 7018 focal and 4570 reticular adhesions). (D) Quantification of immuno-labeling intensities for vinculin and integrin β5 per adhesion in U2OS cells treated for 1 h with DMSO (blue) or CytoD (red) starting 2 h after attachment to VN (see images in Supplementary Figure 3A, B). CytoD significantly reduces vinculin densities in adhesions, but increases β5 densities (Mann-Whitney U test, *** = p < 0.001, 2533 adhesions from 22 DMSO-treated cells and 1410 adhesions from 18 CytoD-treated cells. (E) Quantification of a cell-VN surface attachment assay using CS1-wt (lacking VN receptors including integrin αVβ5) and CS1-β5 (expressing αVβ5) melanoma cells in the presence or absence of: 20 μM CytoD (i.e. only reticular adhesions provide attachment); and/or cyclic RAD peptides (cRAD, non-inhibitory); and/or cyclic RGD peptides (cRGD, inhibit αVβ5-VN ligation). Cell attachment numbers re-scaled relative to: maximum (= 100) CS1-β5 cells in the presence of cRAD peptides, and; minimum (= 0) CS1-β5cells in the presence of CytoD and cRGD peptide. Error bars indicate mean of 6 replicates +/-standard error of the mean (SEM). (F-G) Representative confocal images of talin 1-null mouse embryonic stem cells (mES talin 1 -/-) plated for 3 h on 10 μg/ml VN and immuno-labeled against integrin β5 and talin (‘53.8’ antibody), 48 h after transfection with either a control siRNA (F) or Talin 2-specific siRNA (G). (H) Quantification of cell spread area following control (CTRL) talin 2-targeting oligonucleotide treatments based on experiments described in Figure 3 F-G. (I) Single cell imaging-based quantification of mean integrin β5 intensity in segmented adhesions, standardized as fold change relative to control (CTRL). 20-40 cells imaged per condition, per replicate in H, I. Error bars indicate means of 3 replicates +/-SEM. Scale bars: 10 μm.

To assess the role of talin in reticular AC formation, talin-1-null mouse embryonic stem cells (mES talin-1 -/-) were transfected with talin-2-targeting siRNA. Reduction of talin caused reduced cell spreading (Fig.3F-H; Supplementary Fig.3F-G) (Zhang et al., 2008) and ablated focal ACs (Fig.3F-G). However, integrin β5 was more densely concentrated within reticular ACs (Fig.3F,G,I), similar to cells treated with cytochalasin D (Fig.3A-D).

Integrin β3 and β5-negative CS1-wt cells do not attach to vitronectin, while CS1 cells stably expressing β5 (CS1-β5) attached strongly to VN (Fig. 3E) and formed both focal and reticular ACs. CS1-β5 cells treated with cytochalasin D attached approximately half as strongly as unperturbed CS1-β5 cells, demonstrating that reticular adhesions are able to facilitate cell attachment in the absence of F-actin. This residual adhesion was blocked by competitive inhibition of αVβ5-VN binding using cyclic RGD peptides (D’Souza et al., 1991), confirming αVβ5 specificity (Fig. 3E). Thus, reticular ACs form in the absence of both F-actin and talin and are able to facilitate attachment in the absence of focal adhesions.

### Reticular AC composition is unique

Immunofluorescence screening did not identify any consensus adhesome components present in reticular ACs (Fig. 1; Supplementary Fig.2). We therefore used mass spectrometry to determine their composition. U2OS cells were treated with either DMSO or cytochalasin D, followed by ventral membrane complex isolation and processing for mass spectrometry. 199 proteins were identified in the control condition, 18 of which were consensus adhesome components (Fig.4A) (Horton et al., 2015). Conversely, cytochalasin D-treated samples revealed 53 proteins selectively associated with reticular ACs, only one of which was a consensus adhesome protein (tensin-3). Four of these proteins were discounted from further analysis due to exceptionally high representation in the CRAPome database (Mellacheruvu et al., 2013), leaving a reticular adhesome of just 49 proteins. Lower diversity in the reticular adhesome supports evidence of relative homogeneity in both integrin clustering density (Fig.2D) and integrin dynamics (Fig.2P), indicating that reticular ACs are more homogeneous, including across their lifespan. Gene ontology analysis of the reticular adhesome revealed enrichment of terms relating to membrane organisation and endocytosis (Fig.4B-C), consistent with the mass spectrometric identification of a number of known endocytic adaptors. Six of these were validated by immunofluorescence, including ITSN1 (Fig.4D), NUMB (Fig.4E), WASL, DAB2, EPS15L1 and HIP1R (Supplementary Fig.4C-F).

**Figure 4.**
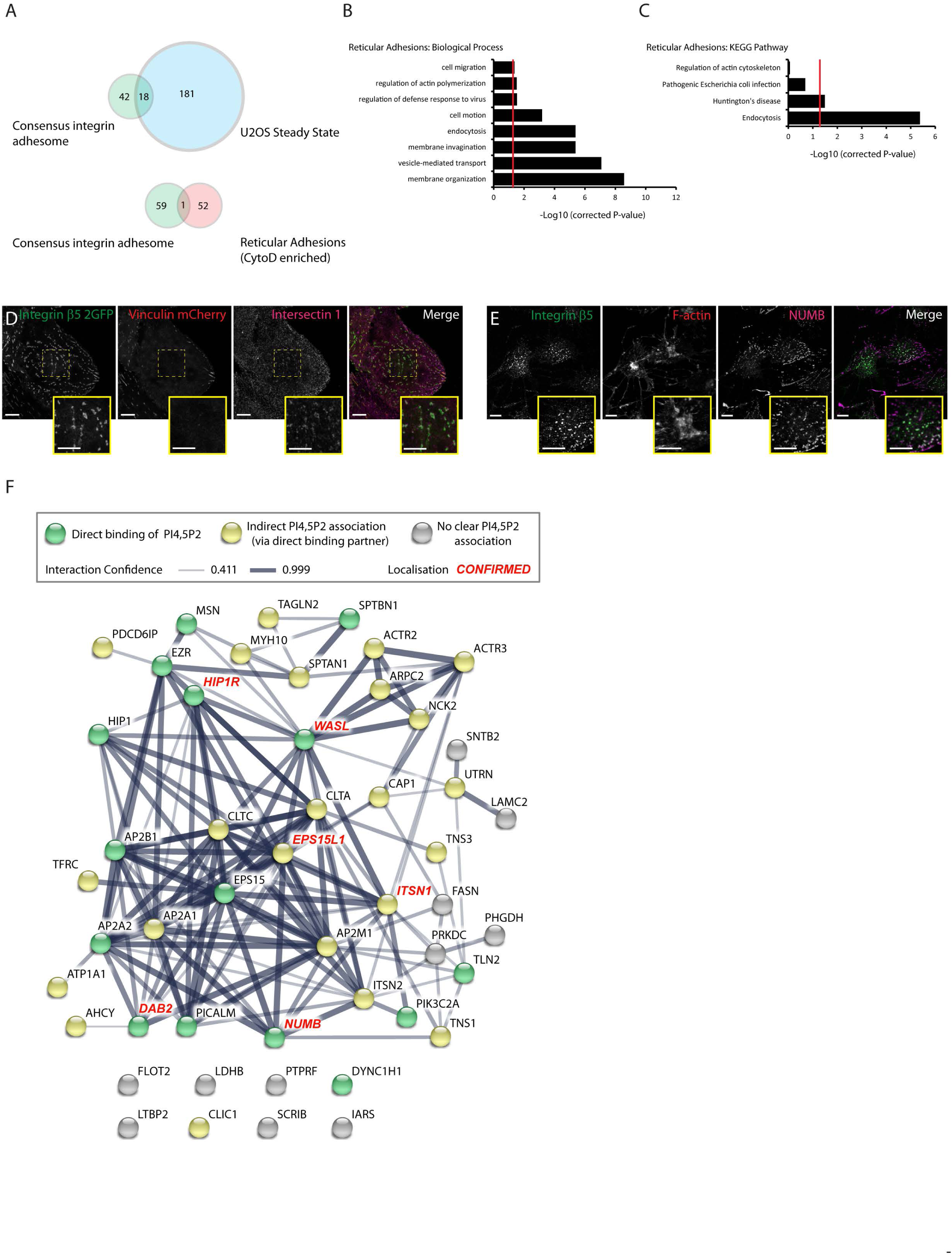
Mass spectrometry reveals the distinct reticular adhesome. (A) U2OS cells were plated in complete medium for 3 d on tissue culture plastic then treated with either DMSO (steady state) or CytoD (reticular enriched) and mass spec analysis was performed of the remaining adhesions after cell removal. Venn diagrams of proteins identified by Mass spec analysis of ventral membrane preparations isolated from each condition overlayed with the 60 consensus fibronectin-adhesome proteins (Horton., 2015) are presented. (B,C) Gene-ontology analysis of reticular adhesion enriched proteins (CytoD-treated) showing terms from Biological Process (B) and KEGG pathway analysis (C) significantly enriched over whole cell proteome. (D) Confocal images of U2OS β5V cells attached to 10 μg/ml vitronectin (VN)-coated surfaces and immuno-labeled against intersectin 1. (E) Confocal images of U2OS cells cultured on glass coverslips for 72 h then treated with 20 μM CytoD for 2 h and immuno-labeled against integrin β5 and NUMB along with staining of F-actin. (F) STRING interaction network of reticular adhesion enriched proteins.

### The balance between reticular and focal ACs is shaped by PIP status

The putative protein interaction network for the identified reticular AC proteins was extremely dense, with many components reported to bind phosphatidylinositol 4,5-bisphosphate (PI-4,5-P2; Fig.4F; Supplementary Table 1). This suggested that reticular ACs may be sensitive to phosphatidylinositol (PIP) signalling. We therefore performed a small RNAi screen to determine whether PIP regulators influence the ratio of focal to reticular adhesions (Fig.5A-B; Supplementary Figure 5A-E). In five out of six cases where PIP regulator depletion would be expected to reduce PI-4,5-P2 levels (PI4KA, PI4K2B, PIP5K1B, PIP5K1C and PTEN), a shift in β5-2GFP intensity ratio was observed from reticular to focal ACs. Correspondingly, depletion of PIK3C2A, which generates phosphatidylinositol-3,4,5-trisphosphate (PIP3) from PI-4,5-P2, caused a relative shift from focal to reticular ACs. To understand how β5-2GFP intensity ratios reflect changes specific to reticular and/or focal ACs, intensities in each AC type were independently assessed (Fig.5C). Depletion of targets that normally produce PI-4,5-P2 reduced β5-2GFP levels in both AC types. Yet because effects were more pronounced for reticular ACs, the ratio to focal ACs decreased. In contrast, PIK3C2A depletion (to reduce PIP3 levels) perturbed only focal ACs, thus causing an increase in reticular to focal intensity ratios.

**Figure 5.**
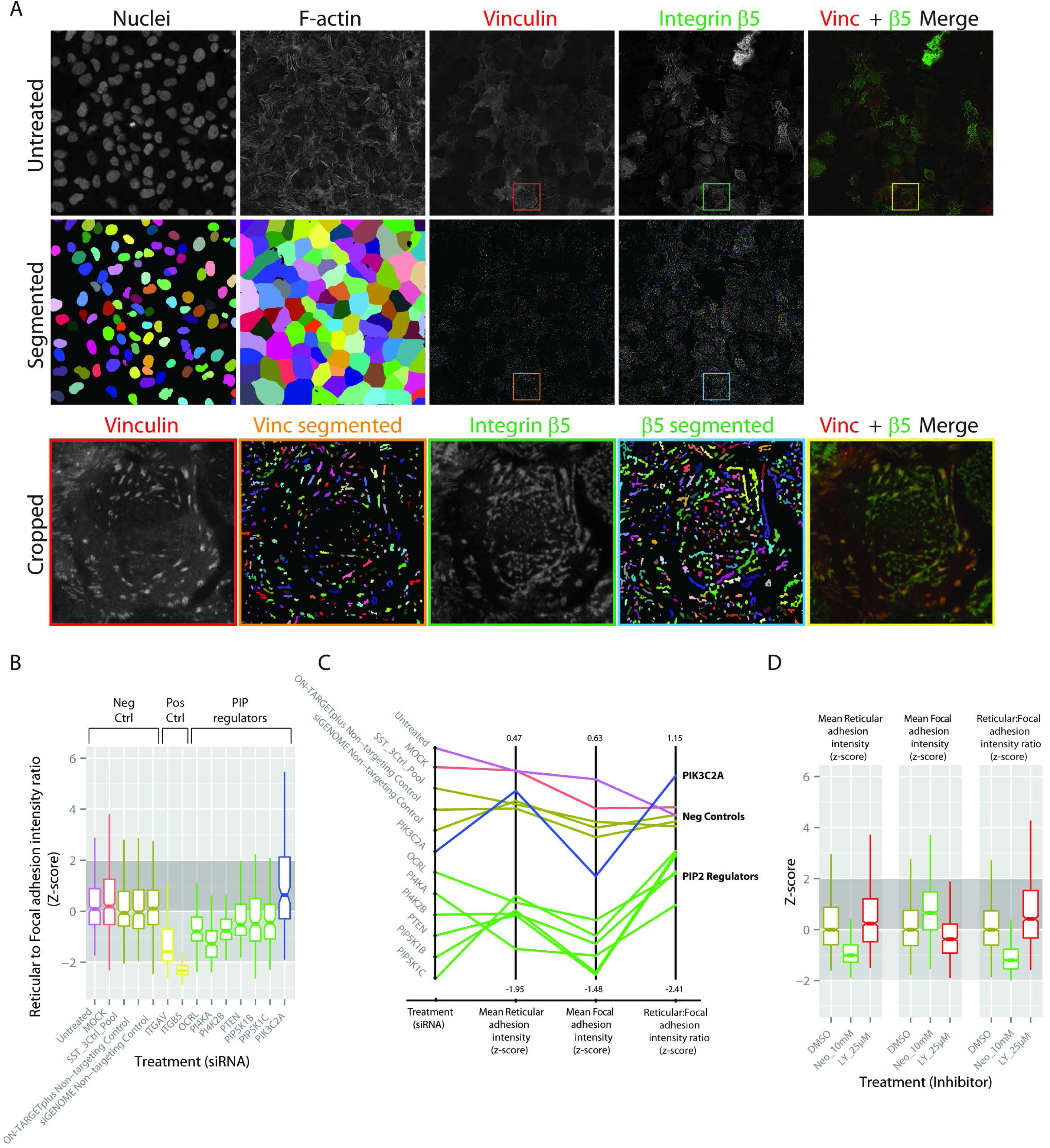
Reticular versus focal adhesion balance is shaped by PIP status. (A) Representative images illustrating segmentation of cells based on DAPI and F-actin staining and subsequent segmentation of vinculin mCherry and integrin β5-2GFP fluorescence (shown for an individual cell; cropped). (B) Quantification of integrin β5 intensity ratios (represented as Z-scores) between reticular and focal adhesions following knockdown of PIP2 and PIP3 regulators. (C) Quantification of mean focal and reticular adhesion integrin β5 intensity following knockdown of PIP2 and PIP3 regulators. (D) Quantification of mean integrin β5 intensity focal and reticular adhesion intensity and ratio following 30 min treatment with 10 mM Neomycin (PIP2 binding inhibition) or 25 μM LY294002 (inhibition of PIP3 generation).

We validated these findings through inhibition of PI-4,5-P2 binding (neomycin) or PIP3 formation (LY294002) (Fig.5D; Supplementary Fig.5F-H). Neomycin reduced β5-2GFP intensities in reticular ACs while increasing intensities in focal ACs, shifting overall ratios from reticular to focal adhesions. Conversely, LY294002 caused increased reticular AC intensities and reduced focal adhesion intensities, thereby raising reticular to focal β5-2GFP intensity ratios. Thus, both siRNA and inhibitor treatments indicated that focal and reticular ACs are in an equilibrium, within which PI-4,5-P2 promotes reticular ACs, while PIP3 promotes focal ACs.

### Reticular ACs persist throughout cell division when focal ACs disassemble

siRNA-mediated knock-down of integrin β5 reduced cell proliferation (Fig.6A), but without affecting S-phase progression (Fig.6B). We therefore analysed the potential role for β5 in mitosis. Remarkably, unlike classical ACs, integrin β5-containing reticular ACs persisted throughout cell division (Fig.6C-I, Supplementary Movie 6) and remained free of consensus adhesome components as well as of integrin β3 (Supplementary Fig.6A-F). This suggests a selective role for αVβ5 in mitotic cell attachment.

**Figure 6.**
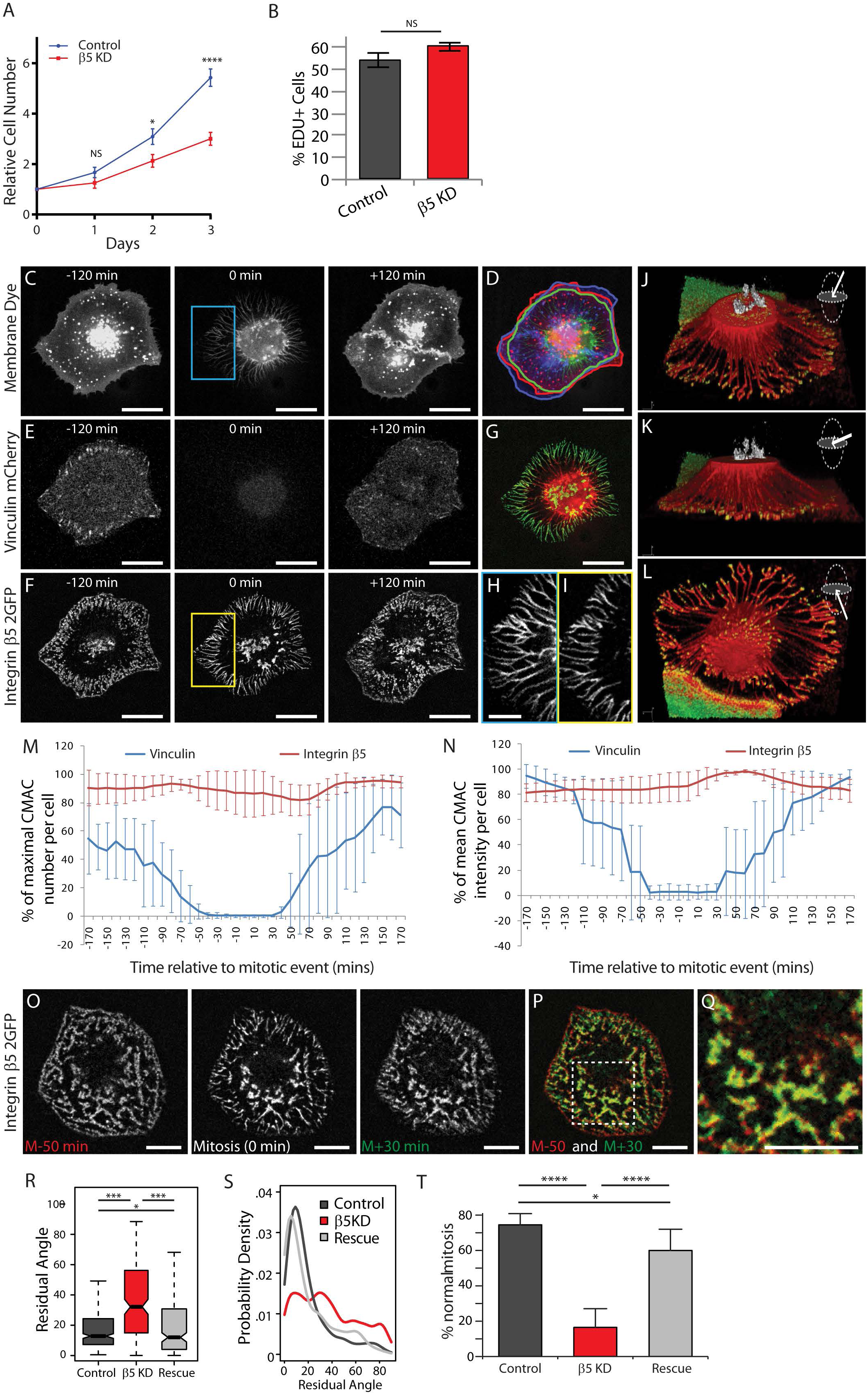
Reticular adhesions persist during mitosis and transmit spatial memory from pre-mitotic to post-mitotic daughter cells. Control or integrin β5 knockdown U2OS cells were assessed for proliferation defects by determining a growth curve over 3 days post attachment (A) or by counting the percentage of EDU-positive cells (B). U2OS cells labeled with a far-red membrane dye (C) as well as stably expressing vinculin mCherry (E) and integrin β5-2GFP (F) were replated on 10 μg/ml vitronectin and imaged every 10 min at the ventral cell-substrate interface via spinning-disc confocal microscopy during mitosis (see **Supplementary Movie 6**). Selected images from each channel show a cell 120 min before, during and 120 min after mitosis. An overlay of membrane labeling (with cell boundaries outlined) at these time points (D, −120 min = red, 0 min = green, +120 min = blue) highlights recovery of the premitotic adhesion footprint by the post-mitotic daughter cells. Membranous retraction filaments formed during mitosis (H; cropped from blue ROI in C) overlap exactly with integrin β5-2GFP-positive adhesion complexes (I; cropped from yellow ROI in F). Scale bars C-F = 10 μm’ H and I = 5 μm. (J-K; see **Supplementary Movie 7**) Three alternate views (above (J), beside (K) and below (L); orientation indicated by arrows in planar schematics) of a 3D confocal-reconstructed mitotic cell showing condensed DNA (white), cell membrane labelling (red; cut through to expose DNA) and integrin β5-2GFP labelling of reticular adhesions (green). Note retraction fibre attachment at sites of integrin β5-2GFP labelling. (M-N) Quantification of vinculin-positive adhesion complex number (M, blue) vs integrin β5-2GFP-positive adhesion complex number (M, red) reveals consistent sustainment of integrin β5-positive adhesion complexes during mitosis, and the complete loss of vinculin-positive adhesion complexes. Similarly, quantification of integrin β5-2GFP clustering density (N, red) indicates no decrease in integrin β5 clustering density, while vinculin mCherry clustering density (N, blue) consistently falls to zero during mitosis before post-mitotic recovery. Error bars in M and N = mean of adhesion values from 5 cells +/-95% CI. (O) Confocal images of integrin β5-2GFP-labeled adhesions in a U2OS cell at three time points relative to mitosis (-50 min (pre); 0 min, +30 min (post)). (P) Overlay of pre and post mitosis adhesions, cropped and zoomed in O, confirm the persistence and morphological consistency of reticular adhesions throughout mitosis (see **Supplementary Movie 9**). (R-T) Based on data exemplified in **Supplementary Figure 6** and **Supplementary Movies 10-14**. (R-S) Quantification relative to control siRNA (Control; n = 297 cells) of integrin β5 knock down (β5 KD; n = 176 cells) and β5 rescue (Rescue; n = 195 cells) effects on spatial memory transmission between Hela cell generations, defined by residual angle measurement between the pre-mitotic cell major axis and the cell division axis. Box-plots (R) and probability density plots (S) indicate the distribution of residual angles. Significant differences indicated (Mann-Whitney U test, * = p < 0.05, *** = p < 0.001). (T) Quantification of the percentage of rounding cells that progressed through normal mitosis; mean of three separate experiments +/-SD. Significant differences indicated (two-tailed T test, * = p < 0.05, *** = p < 0.001).

The pre-mitotic footprint of the mother cell is often transmitted with high precision to post-mitotic daughter cells (Fig.6D) (Mali et al., 2010; Toyoshima and Nishida, 2007). During the rounding phase of mitosis, this footprint was visibly demarcated by both membrane dye-labeled retraction fibres and integrin β5-2GFP-labeled reticular ACs (Fig.6G). Indeed, the exquisite correspondence between retraction fibres (Fig.6H) and reticular ACs (Fig.6I) was highlighted by confocal-based threedimensional visualisation of a similarly staged mitotic cell (Fig.6J-L; Supplementary Movie 7). Here, retraction fibres angled down and attached precisely at sites decorated with β5-labeled reticular ACs. This strongly suggests a role for reticular ACs in mediating cell-ECM attachment during cell division and, consequently, in the transmission of inter-generational spatial memory. Supplementary Movie 8 exemplifies the dynamic re-spreading of post-mitotic daughter cells to cover the reticular adhesion-defined retraction fibre footprint of the pre-mitotic mother cell. Thus, reticular adhesions appear to actively mediate the transmission of spatial memory between cell generations.

Quantitative analyses confirmed the observations above, showing that the number as well as intensity of vinculin-positive (focal) ACs fell to virtually zero during mitosis, while β5-positive AC numbers and β5 intensity were maintained (Fig.6M-N). Notably, detailed comparison of reticular ACs before, during and after mitosis (Fig.6O-Q and Supplementary Movie 9) confirmed that, particularly in central reticular ACs, the geometry of macromolecular protein complexes remained virtually unchanged from one cell generation to the next. In contrast, peripheral reticular ACs (generally associated with mitotic retraction fibres) underwent remodelling characterised by both narrowing and intensification of the complex. At STORM resolution, central mitotic reticular ACs were indistinguishable in organisation from interphase reticular ACs (Fig.2X-Z, Supplementary Fig.6I). In contrast, peripheral mitotic retraction fibre-associated reticular ACs were highly linearised and condensed, as confirmed by quantification of nanocluster nearest neighbour distances and molecular localisation counts per nanocluster (Supplementary Fig.6I-K). Such molecular-scale remodelling functionally implicates reticular ACs in the mechanical process of cell-ECM attachment during cell division.

### Reticular ACs are required for cell division and inter-generational spatial memory-transmission

Because integrin β5-positive reticular ACs persisted in dividing cells, we tested reticular AC function via RNAi-mediated depletion of β5 in HeLa cells, given both their expression of reticular ACs (Supplementary Fig.1B) and their extensive characterisation during cell division (Held et al., 2010). In addition to recovery of the pre-mitotic footprint by daughter cells (detailed above), transmission of spatial memory from mother to daughter cells is also reflected in the preference for cells to divide along the major axis of the pre-mitotic mother cell. This determines the spatial arrangement of daughter cells (Jime et al., 2007; Minc et al., 2011). We therefore measured the residual angle between the pre-mitotic major axis and the mitotic division axis (Fig. 6R-S). Residual angle distributions were strongly skewed towards zero (indicating spatial memory retention) in control cells and those in which β5-depletion had been rescued. Contrastingly, mitotic axis orientation in integrin-β5-depleted cells was almost random relative to the pre-mitotic major axis, indicating a loss of spatial memory. Thus, reticular ACs are indeed critical for inter-generational spatial memory transmission during cell division.

While approximately 75% of control cells underwent normal cell division, only around 15% of β5-depleted cells completed normal cell division (Fig.6T). Indeed, compared to controls (Supplementary Fig.6L; Supplementary Movie 10), several distinct cell division defects were observed in β5-depleted cells. Defects included significantly delayed mitosis often with incomplete cytokinesis, repeated cell rounding and re-spreading without division, and failure of cytokinesis resulting in bi-nucleate daughter cells (Supplementary Fig.6M; Supplementary Movies 11-13). The frequency of such errors was greatly reduced by β5-EGFP rescue (Fig 6T; Supplementary Fig.6N-O; Supplementary Movie 14). Together, these findings demonstrate that integrin β5-mediated reticular ACs are essential for the normal progression of cell division in cultured HeLa cells.

## Discussion

Our major finding is the identification and characterization of a novel cellular structure, the reticular adhesion. This new class of cell-ECM adhesion complex is the first with a clear role in mediating cell-ECM attachment during cell division. Reticular adhesions form in a diverse array of cell types and are characterised by both the presence of integrin αVβ5 and the absence of consensus adhesome components. Reticular adhesions are unique in that they are functionally independent of both F-actin and talin, yet they also display distinct morphology and dynamics, as well as differential regulation by PIP signalling relative to classical adhesions.

Remarkably, reticular adhesions have remained unrecognized despite 30 years of intensive integrin research and early studies displaying similar reticular αVβ5 labeling patterns (Wayner et al., 1991). Instead, the literature is dominated by studies of adhesion complexes formed within the first hours of cell attachment, when a pattern of temporal evolution is well established; early nascent adhesions and focal complexes mature towards focal and fibrillar adhesions. Such attachment and spreading onto new surfaces is analogous to initial cell responses after tissue wounding, when membrane protrusion and adhesion complex turnover drive engagement with new microenvironments, enabling migration and repair. While our current understanding of cell-matrix interactions largely derives from such dynamic contexts, the discovery of reticular adhesions now demands equivalent analyses of stable cell adhesion *in situ*.

Reticular adhesions persist throughout mitosis and provide a solution to the paradox of mitotic cell-ECM adhesion, when all previously known ACs do and must disassemble for effective mitosis and yet cells typically remain anchored (Dao et al., 2009; Ferreira et al., 2013; Maddox and Burridge, 2003; Ramkumar and Baum, 2016). In addition to enabling mitosis itself, cell-ECM attachment is also crucial for spatial memory transmission between cell generations, including defining the axis of cell division (Jime et al., 2007; Minc et al., 2011). To date it has been unclear how residual adhesion is maintained during mitosis, nor how re-spreading is guided thereafter. Reticular adhesions now provide mechanisms underpinning both phenomena. Interestingly, the unique characteristics of reticular adhesions appear precisely suited to these roles in cell division. For instance, F-actin independence decouples reticular adhesions from the massive cytoskeletal remodelling that occurs during cell rounding. Importantly, a key role for reticular adhesions in cell division is confirmed by integrin β5 depletion, which caused multiple mitotic defects and disturbed spatial memory transmission between cell generations.

Whilst we find an important role for αVβ5 in mitosis and proliferation, cells can also divide on extracellular matrix ligands not engaging αVβ5, such as laminin. This implies that, when required to, cells can deploy alternate adhesion receptors for mitotic anchorage. Whether other integrins can play a similar mitotic role to αVβ5 in cells plated on different matrix proteins should be the subject of further studies. However, β5 knockout mice have a relatively limited phenotype, suggesting redundancy of function for different integrin sub-families and/or a specialised role for αVβ5 in regulating division within specific matrix environments. Nonetheless, both β5 knockout (Lane et al., 2005) and overexpression (Coudert et al., 2014) in mice cause deficiencies in osteoblast/osteoclast function and differentiation, potentially reflecting mitotic defects in cells on flat, rigid, RGD-rich substrates. Our findings now suggest that such defects might include spatial memory transmission failures including loss of appropriate division axis-alignment; a known cause of differentiation errors (Akanuma et al., 2016). Indeed, flat, rigid, RGD-rich *in vivo* environments may be somewhat analogous to long-term cell culture conditions, where we show αVβ5-mediated reticular ACs to be the dominant adhesion type. The fact that αVβ5 is selectively utilized in long-term culture suggests that αVβ5 provides a mechanistic advantage during cell division in such contexts, potentially reflecting the contribution of αVβ5 to spatial memory. Intriguingly, αVβ5 is expressed at high levels in a number of proliferative diseases, raising the possibility that heightened β5 expression may promote disease progression by enhancing cell division within specific matrix environments. It is thus now important to develop an understanding of integrin αVβ5 and reticular adhesion status in 3D *in vivo* environments, including within both physiological and disease settings. One indication to this end may be a focal adhesion-independent role for αVβ5 in 3D skin formation and tumour invasion (Duperret et al., 2015).

Reticular adhesions lack not only F-actin, but virtually all consensus adhesome components. Most notably, both talin and kindlin are absent, despite being considered necessary and ubiquitous activators of integrins, as well as key mechanosensory machineries (Klapholz and Brown, 2017). Moreover, perturbation of talin and F-actin indicate the functional independence of reticular ACs from these normally pivotal proteins. Our proteomic analysis instead identified a distinct reticular adhesion adhesome, highly enriched in PI-4,5-P2-binding proteins. These include clathrin-mediated endocytosis machineries, such as Dab2 and Numb, previously shown to interact directly with the integrin β5 cytoplasmic tail *in vitro* (Calderwood et al., 2003). These data are consistent with recent evidence of integrin-mediated ECM attachment to both 3D substrates and low-tension environments via clathrin-coated structures (CCSs) (Elkhatib et al., 2017; Leyton-Puig et al., 2017). Given that both reticular adhesions and CCSs can form in the absence of talin, it follows that some integrins may not depend on talin for their activation. In this context, it is also notable that we observe near identical nanoscale integrin β5 clustering between talin-positive and talin-negative (reticular) adhesions during interphase, despite previous suggestions that talin determines nanoscale integrin organisation (Liu et al., 2015). Thus, both in terms of integrin activation and organisation, it is possible that either alternative proteins can replace talin functions in reticular adhesions, or that β5 ligand-binding and nanoscale organisation are independent of cytosolic regulators. Regardless, the composition, regulation and function of integrin-mediated adhesion complexes appears more diverse than previously recognised.

In conclusion, we here define reticular adhesions, a novel cellular structure and new AC class. Functionally, by mediating cell-ECM attachment during mitosis, reticular adhesions provide the first known solution to the paradox of mitotic cell attachment, when classical ACs must disassemble but cells must also remain ECM-attached. These discoveries not only delineate a new and specific form of adhesion with a ground-breaking functional role, but also highlight important and somewhat overlooked aspects of adhesion biology that now merit further attention in future research.

## Methods

### Cell culture, plasmid generation, transfection and stable cell generation

#### Cell Culture

U2OS human osteosarcoma cells (ATCC), Hela human cervical carcinoma cells (ECACC), MCF-7 human breast carcinoma cells (ATCC), A549 human lung carcinoma cells (ECACC) and A375 human melanoma cells (ECACC) were maintained in DMEM (Gibco) supplemented with 10% FBS (Sigma) and 2mM L-glutamine (Gibco). U2OS β5V cells stably expressing integrin β5-2GFP and vinculin mCherry were maintained with the addition of 600 μg/ml Geneticin (G-418 sulphate; Gibco). H1299 human non-small lung cancer cells (kind gift from Benny Geiger, The Weizmann Institute of Science, Rehovot, Israel) and CS-1 wild-type hamster melanoma cells were cultured in RPMI-1,640 (Gibco) medium supplemented with 10% FBS and 5 mg/ml L-glutamine. CS-1 cells stably expressing integrin β5 (CS1-β5) were maintained with the addition of 500 μg/ml G-418. BT549 (ductal breast carcinoma, ATCC) cells were maintained in RPMI 1640 medium containing 10% FBS and 1 mM L-glutamine. Mouse aortic endothelial (MAE) cells (ATCC) were grown in RPMI 1640 medium with 5% FBS. All live cells were incubated and imaged in a humidified environment at 37°C with 5% CO2.

#### DNA plasmid generation and sourcing

For construction of integrin β5-2GFP, EGFP was duplicated in a pEGFP-N1 backbone vector (gift of Dr Pat Caswell, University of Manchester, UK), then a full-length integrin β5 cDNA (kindly provided by Dr Errki Ruoslahti, Burnham Institute) was subcloned into the 2XEGFP-N1 vector using the EcoRI site of the original pEGFP-N1 vector. The vinculin mCherry plasmid was kindly provided by Dr Vic Small (IMBA, Austria). Csk-GFP was kindly provided by Dr Akira Imamoto (University of Chicago, USA).

#### Transfection and stable cell line generation

Cells were transfected at confluences ranging from 70–90%, 24 h after plating into 12 well culture plates (except where otherwise stated). For DNA plasmid transfection, 0.3-2 μg of total DNA was mixed with 0.5-3 μ! of Lipofectamine Plus or Lipofectamine 2000 (Thermo Fisher Scientific) according to the manufacturer’s instructions. For RNA transfection, except where otherwise stated, 15-30 pmol of siRNA was transfected together with 0.5-3 μl of RNAiMAX (Thermo Fisher Scientific). Cells were typically imaged 24 to 48 h after transfection. U2OS β5V cells expressing integrin β5-2GFP and vinculin-mCherry were established via manual single colony selection followed by selection with 600 μg/ml G-418.

#### ECM surface coating

Cells were typically assayed in 96-well glass-bottomed plates (0.17 mm optical glass; Matrical Bioscience). Glass coating was performed at 37°C for 2 h after blocking with 1% heat-denatured bovine serum albumin (BSA; Sigma-Aldrich) for 2 h at 37°C. ECM ligand coating concentrations were 10 μg/ml except where otherwise indicated (where vitronectin concentrations were varied). Vitronectin and Fibronectin were purified from human plasma as detailed previously (Smilenov et al., 1992; Yatohgo et al., 1988), while purified Laminin was acquired commercially (Sigma-Aldrich).

### Antibodies, immunofluorescence labelling and immuno-blotting

Primary antibodies used for immunofluorescence and/or immuno-blotting include: anti-integrin β5 (15F11; MAB2019Z) (Millipore); anti-integrin αVβ5 (P1F6) (Abcam); polyclonal (rabbit) anti-integrin β5 (ab15459) (Abcam); anti-integrin β5 (4708S) (Cell Signalling Technology); anti-integrin αv (LM142) (Merck Millipore); anti-pan-talin (1 and 2) (53.8) (BioRad); anti-talin 1 (TA205) (Santa Cruz); anti-talin 2 (68E7) (Abcam); anti-integrin αVβ3 (LM609) (Abcam); anti-integrin β3 (AP3) (Abcam); anti-integrin β1 (LM534) (Millipore); anti-vinculin (hVIN-1) (Sigma Aldrich); anti-vinculin (V9131) (Sigma-Aldrich); anti-intersectin 1 (HPA018007) (Atlas Antibodies, Sigma-Aldrich); anti-NUMB (2733) (Cell Signaling Technologies); anti-EPS15L1 (HPA055309) (Atlas Antibodies, Sigma-Aldrich); anti-HIP1 (HPA013606) (Atlas Antibodies, Sigma-Aldrich); anti-WASL (HPA005750) (Atlas Antibodies, Sigma-Aldrich); anti-DAB2 (12906) (Cell Signaling Technologies); anti-paxillin (5H11) (Sigma Aldrich); anti-FAK (BD Biosciences); anti-zyxin (H-200) (Santa Cruz); anti-kindlin 2 (ab74030) (Abcam); anti-ICAP1 (115228) (Abcam); anti-DOK1 (HPA048561) (Atlas Antibodies, Sigma-Aldrich); polyclonal (rabbit) anti-phosphotyrosine (1000) (Cell Signaling); anti-cytokeratin (27988) (Abcam); anti-beta tubulin (DM1A) (Thermo Fisher Scientific); anti-vimentin (8978) (Abcam). Anti-mouse and anti-rabbit secondary antibodies conjugated with Alexa 488, 568 or 647 were used as appropriate (Thermo Fisher Scientific). For fixed F-actin labelling, phalloidin pre-conjugated with Alexa 488, 568 or 647 was used as appropriate (Thermo Fisher Scientific). DAPI (4’, 6-Diamidino-2-Phenylindole, Dihydrochloride; Thermo Fisher Scientific) nucleic acid stain was used as a nuclear marker as appropriate.

Immunofluorescence labeling was performed either manually or using liquid-handling robotics (Freedom EVO, Tecan) to minimise experimental variability, as described previously (Kiss et al., 2015). In either case, standardised procedures were used except where otherwise stated. Cells were fixed with 4% paraformaldehyde (PFA; Sigma-Aldrich) for 20 min, washed 3× with PBS and permeabilised using 0.1% TX-100 (Sigma-Aldrich) for 5 min at room temperature. Cells were then blocked for 15 min with 1% BSA in phosphate-buffered saline (PBS) (PBS/BSA). Primary antibody immuno-labelling then proceeded at room temperature for 30 min. After further PBS/BSA washing, secondary antibodies conjugated with either Alexa 488, 568 or 647 fluorophores (used as appropriate) were then applied for 30 min at room temperature. Finally, cells were washed with PBS.

Immuno-blotting was performed on SDS-polacrylamide gels with proteins transferred to Immobilon-P-Membranes (Millipore). Membranes were probed with anti-talin 2 mouse monoclonal (68E7) (Abcam) at 1:10 000 dilution, anti-vinculin (V9131) (Sigma-Aldrich) at 1:150 000 dilution, or anti-integrin β5 (4708S) (Cell Signalling Technology) at 1:1000 dilution. Proteins were detected using the enhanced chemiluminescence reagent (ECL, Amersham Pharmacia Biotech).

### Imaging

Live and fixed cell imaging was primarily performed using a Nikon Ti2-mounted A1R confocal microscope running NIS elements software (Nikon) with a PlanApo VC 60X / 1.4 NA oil-immersion objective. Live cell fluorescence imaging during cell division was also performed using a Yokagoawa CSU-X1 spinning-disk confocal coupled to an Andor EM-CCD, enabling imaging with reduced light exposure. TIRF imaging was performed on a Nikon Ti2 inverted microscope with laser angle optimised to achieve minimal (~90 nm) evanescence wave penetration. For live cell imaging, images were typically acquired at between 0.5 - 5 min intervals for 1 - 8 h, with pixel resolutions between 0.13 - 0.21 μm. During live cell imaging, cells were maintained in normal culture medium, absent FCS / FBS, at 37°C and 5% CO2. Live cell interference reflection microscopy (IRM) was performed on a Zeiss LSM 510 confocal microscope via a Plan-Apochromat 63X /1.4 NA oil objective, with post-sample dichroic mirror displacement allowing reflected (561 nm) laser light to reach the detector.

Fluorescence recovery after photobleaching (FRAP) analyses were performed via confocal and analysed as described previously (Coló et al., 2012; Li et al., 2010). Briefly, three sequential images were acquired of integrin β5-2GFP and vinculin mCherry in U2OS β5V cells prior to bleaching, enabling robust recovery standardization. Both reticular and focal adhesions (2-3 each per cell) were then bleached using 35% of maximal 488 nm laser power over 40 rapid iterations (< 3 s per cell). Recovery was monitored for a total of 1875 seconds, with intervals of 6 s for the first 120 s (to optimally capture early rapid early recovery dynamics), and intervals of 45 s thereafter (to minimise impacts of monitoring).

Stochastic optical reconstruction microscopy (STORM) was performed in U2OS cells fixed during either interphase or mitosis. Cells were labeled using the rabbit-derived polyclonal anti-integrin β5 (ab15459) and with AlexaFluor 405-AlexaFluor 647 double labeled secondary. Classical adhesions were demarcated by vinculin mCherry, allowing definition of adhesion type from overlaid diffraction-limited images. Secondary antibodies (Jackson ImmunoResearch) were labelled in-house, as previously described (Bates et al., 2007). A Nikon N-STORM system with Apo internal reflection fluorescence 100X / 1.49 NA objective was used, with images acquired via EM-CCD camera. Prior to STORM imaging, TIRF images were acquired to enable standard (diffraction-limited) definition of reticular and focal adhesions using the criteria defined above. Thereafter, 647 nm laser light was used to excite Alexa 647, with 405 nm light used for reactivation. Standard STORM imaging buffer was used, containing 100 mM Cysteamine MEA, 0.5 mg/ml glucose oxidase, 40 μg/ml catalase, and 5% Glucose (all Sigma Aldrich).

### Image Analysis

Patch Morphology Analysis Dynamic software (Digital Cell Imaging Laboratories, Belgium) was used for analysis of static (fixed) and dynamic (live) cell imaging data, except where otherwise specified. Analysis strategy and parameterisation are as described previously (Hernández-Varas et al., 2015; Kiss et al., 2015; Kowalewski et al., 2015; Lock et al., 2014; Shafqat-Abbasi et al., 2016). Briefly, both cells and intracellular adhesion cohorts were segmented according to pixel intensity gradient analysis. A variety of morphological, pixel intensity and dynamic properties were then extracted, for each cell and for each adhesion (Lock et al., 2014). Relationships between each adhesion and its (parent) cell were automatically maintained. Minimal adhesion size was set to 0.3 μm^2^. For live cell data, adhesion tracking parameters included: linear motion interpolation over maximum 1 missing time point; 3 μm maximum adhesion step-size per time point; 4 time point minimum track lifetime. When quantifying differences between reticular and focal adhesions, we used the absence or presence (respectively) of canonical adhesome components as a defining indicator. Specifically, we applied a threshold such that segmented adhesions (delineated by integrin β5) were defined as reticular if they contained less than the mean of background fluorescence values (pixel intensities inside the cell boundary but outside segmented adhesions) plus two standard deviations for a canonical adhesion marker (typically vinculin or talin). Integrin β5-positive adhesions with greater than this value of fluorescence (for the canonical adhesion marker) were classed as focal adhesions.

For FRAP analyses, PAD software was used to segment integrin β5-2GFP-positive adhesions found in the last (3^rd^) pre-bleach image frame. Focal and reticular adhesions were distinguished based on vinculin-mCherry content, as described above. Identical adhesion boundaries (from pre-bleach frame 3) were then used as fluorescence recovery measurement locations for all subsequent image frames. Adhesions adjudged to move during this period were excluded from further analysis. Integrin β5-2GFP fluorescence recovery curves were first standardised relative to intensity fluctuations (including non-specific photo-bleaching) in non-bleached areas of the cell. Thereafter, intensity values in bleached regions were standardised per adhesion as a percentage of the mean of the three pre-bleached images. The standard deviation of percentage recovery, per time point, was also recorded.

STORM data were analysed using Insight3 software (developed by Bo Huang, University of California, San Francisco). First, localisation coordinates were precisely defined via Gaussian fitting. Next, reticular and focal adhesions were segmented and defined using conventional TIRF images of integrin β5 and vinculin, based on the thresholding criteria detailed above. Clustering was then performed on integrin β5 localisations within each adhesion type, revealing coordinate position and localisation counts for integrin nanoclusters found within each adhesion. DBSCAN was used for clustering (Ester et al., 1996), with epsilon (search radius) set to 10 nm and minimum points (within epsilon radius) set to 3. Nearest neighbour distances between nanoclusters and localisation numbers per cluster were assessed for each adhesion type using R.

Three-dimensional rendering and animation of confocal images was performed using NIS elements software. Additional supplementary movies were prepared in FiJi software (Schindelin et al., 2012).

### Mass Spectrometry analysis of the Reticular Adhesome

Four 10 cm-diameter dishes per condition of U2OS cells were cultured for 48 h to 90% confluency then treated with either DMSO or 20 μM Cytochalasin D (Sigma-Aldrich) for 2 hours. To isolate adhesion complexes, cells were incubated with the membrane permeable cross-linker dimethyl-3, 3′-dithiobispropionimidate (DTBP, Sigma Aldrich; 6 mM, 5 min). DTBP was then quenched using 1 M Tris (pH 8.5, 2 min), after which cells were again washed once using PBS and incubated in PBS at 4°C. Cell bodies were then removed by a combination of cell lysis in RIPA buffer (50 mM Tris-HCl, pH 8.0, 5 mM EDTA, 150 mM NaCl, 1% (w/v) TX-100, 1% (w/v) sodium deoxycholate (DOC), 0.5% (w/v) sodium dodecylsulfate (SDS); 3 min) and a high-pressure water wash (10 s). Protein complexes left bound to the tissue culture dish were washed twice using PBS, recovered by scraping in 200 μl recovery solution (125 mM Tris-HCl, pH 6.8, 1% (w/v) SDS, 15mM DTT), and incubated at 70°C for 10 min. Each sample was subsequently precipitated from solution by addition of four volumes −20°C acetone, incubated for 16 h at −80°C, and resuspended in reducing sample buffer.

For mass spectrometric, samples were separated by SDS-PAGE on a 4-12% SDS Bis-Tris gel (Thermo Fisher), stained for 10 min with Instant Blue (Expedeon), and washed in water overnight at 4 °C. Gel pieces were excised and processed by in-gel tryptic digestion as previously described (Horton et al., 2016). Peptides were analysed by liquid chromatography (LC)-tandem MS using an UltiMate 3000 Rapid Separation LC (RSLC, Dionex Corporation, Sunnyvale, CA, USA) coupled to an Orbitrap Elite mass spectrometer (Thermo Fisher). Peptides were separated on a bridged ethyl hybrid C18 analytical column (250 mm × 75 μm I.D., 1.7 μm particle size, Waters) over a 1 h gradient from 8% to 33% (v/v) ACN in 0.1% (v/v) FA. LC-MS/MS analyses were operated in data-dependent mode to automatically select peptides for fragmentation by collision-induced dissociation (CID). Quantification was performed using Progenesis LC-MS software (Progenesis QI, Nonlinear Dynamics, Newcastle, UK; http://www.nonlinear.com/progenesis/qi-for-proteomics/)) as previously described (Horton et al., 2016). In brief, automatic alignment was used, and the resulting aggregate spectrum filtered to include +1, +2, and +3 charge states only. An .mgf file representing the aggregate spectrum was exported and searched using Mascot (1 missed cleavage, fixed modification: carbamidomethyl [C]; variable modifications: oxidation [M]; peptide tolerance: ± 5 ppm; MS/MS tolerance: ± 0.5 Da), and the resulting .xml file was re-imported to assign peptides to features. Three separate experiments were performed and abundance values for proteins identified in analysis were used to determine which proteins were enriched over 2-fold following treatment with Cytochalasin D. While 53 proteins were detected in original mass spectrometry data, 4 were excluded in further analysis due to high representation in the CRAPome database (Mellacheruvu et al., 2013). The putative reticular adhesome interaction network was constructed using the online STRING proteinprotein interaction database (v10) (Szklarczyk et al., 2015) including experimentally validated interactions only, with a medium interaction confidence score (> 0.4). Even at higher confidence (interaction confidence score > 0.7), this network is extremely dense in terms of internal interactions, with 91 known interactions relative to a randomly expected rate of just 11 (relative to proteome-wide interaction frequencies). Biological process- and KEGG pathway-enrichment analyses were also performed using the STRING database.

### siRNA and drug screening against PIP regulators

U2OS β5V cells were treated with pooled siRNAs (4 siRNAs per target; ON-TARGET SMART Pool plus; Dharmacon) via reverse transfection in the inner 60 wells of 96-well optical glass plates. Each plate contained 5 negative (untreated; mock transfected; 3 non-targeting siRNA controls) and 3 positive targeting controls (against EGFP, integrin αv or integrin β5). The screen was repeated twice, with a third validation assay using 4 siRNAs individually, per target (Dharmacon). siRNA sequences are displayed in Supplementary Table 2. To prepare the siRNA library, 1 μ! of each siRNA pool from 2 μl stock was mixed with 30 μl nuclease-free water and added to 96-well glass-bottom plate wells, before drying at RT. To enable reverse transfection, 30 μl of RNAiMAX was first added to 9 ml of Opti-MEM (Thermo Fisher Scientific). 30 μl of this mixture was added to each well, followed by a 30 min incubation. 90% confluent U2OS β5V cells grown in 75 cm^2^ flasks were trypsinized and resuspended with 30 ml of growth medium. 100 μl of the resulting cell suspension was added to each well, and pipetted 5 times to ensure good cell dispersion. The final concentration of siRNA was 15 nM. Cells were incubated for 48 h before fixation with 4% PFA (15 min) and subsequent permeabilization with 0.2% TX-100 in PBS. Finally, cells were incubated for 1 h with DAPI and Alexa 647-conjugated phalloidin before 3x PBS washing.

Cells were imaged with a Nikon A1R confocal microscope with PlanApo VC 60X /1.4 NA oil-immersion objective. Image settings were identical for all samples and repeats. Montage images were acquired and stitched in NIS-elements software, enabling the high-resolution acquisition of ~100 cells and ~5000 adhesions per condition, per experimental repeat. Image data were quantified and analysed using KNIME software (v3.3.2; KNIME). Individual cells were segmented using Voronoi tessellation based on DAPI (nuclei) and phalloidin (cell body) staining. Integrin β5-positive adhesions were then segmented and split using spot detection and the Wählby method (Wählby et al., 2004), respectively. Focal and reticular adhesions were defined based on vinculin mCherry content as described above. Background-corrected intensity values were extracted per channel, for each adhesion, per cell. Mean integrin β5 intensity values in reticular adhesions were divided by values in focal adhesions, to generate the relative intensity ratio. All values were Z-score standardized using robust statistics (median and median absolute deviation) relative to the combination of (3) nontargeting siRNA controls per 96-well plate. Resulting response distributions were plotted using R software. Boxplot notches indicate +/− 1.58 times the interquartile range divided by the square root of observation number, approximating the 95% confidence interval.

U2OS β5V cells cultured and plated as described above, including 48 h incubation in 96-well optical glass plates, were treated for 30 min with either: DMSO (control); 10 mM Neomycin (PIP2 binding inhibition), or; 25 μM LY294002 (inhibition of PIP3 generation). Cells were then fixed, permeabilised and labeled as described above. Imaging and analysis was again performed using KNIME, as described above for siRNA screening.

### Talin knock-down and response analysis

Talin 1-null mouse embryonic stem cells (mES talin 1 -/-; kind gift of David Critchley, University of Leicester) were transfected using RNAiMAX (Thermo Fisher Scientific) as per the manufacturer’s instructions using either non-targeting control siRNA (ON-TARGETplus Non-targeting Control; 5′-UGGUUUACAUGUCGACUAA-3′) or talin 2-specific siRNAs designated: talin-2 siRNAl (5′-GCAGAAUGCUAUUAAGAAAUU-3) talin-2 siRNA2 (5′-CCGCAAAGCUCUUGGCUGAUU-3′), or; talin-2 siRNA3 (5′-AAGUCAGUAUUACGUUGUUUU-3′). siRNAs were synthesized by GenePharma (Shanghai, China). Cells were incubated for 48 h then plated for 3 h on 10 μg/ml VN. Fixation and permeabilisation conditions were tuned to retain cytoplasmic talin 2, as described previously (Kiss et al., 2015). Briefly, labelling was performed using liquid-handling robotics (Freedom EVO, Tecan) to reduce experimental variability. Cells were fixed with 2% PFA for 10 min, washed with PBS and permeabilised using 0.1% TX-100 for 5 min at room temperature. Cells were then blocked for 15 min with 1% PBS/BSA. Immuno-labeling followed at room temperature for 30 min, targeting integrin β5 (polyclonal Ab; ab15459, Abcam) and talin (pan-talin mouse monoclonal Ab ‘53.8’, BioRad or; anti-talin 2 mouse monoclonal Ab ‘68E7’, Abcam). After washing with 1% PBS/BSA, Alexa 488 and 647 secondary antibodies were applied, targeting rabbit and mouse primary antibodies, respectively. Images of integrin β5and residual talin 2 were acquired with a Nikon A1R confocal using an oil-immersion objective (PlanApo VC 60X / 1.4 NA). Image analysis was performed using PAD software to record residual talin (mean) intensities per cell (displayed as knock down efficiency), mean β5intensities per segmented adhesion (per cell), and cell area. All values were scaled as fold-change relative to control siRNA. 20-40 cells were imaged per condition in each of 4 experimental repeats with talin-2 siRNA1, or single confirmatory experiments with talin-2 siRNA2 and 3. Immunoblotting was performed as described above.

### Integrin β5 knock-down and mitotic analysis

siRNA used for knockdown of β5 targeted the sequence 5’-GGGAUGAGGUGAUCACAUG-3’ and was obtained from Dharmacon. For rescue of β5 expression, an siRNA-resistant WT β5-EGFP clone was generated using the QuickChange IIXL site-directed mutagenesis kit (Agilent Technologies) to introduce silent mutations in the siRNA target sequence. The primers were: forward 5’-AGCCTATGCAGGGACGAAGTTATTACCTGGGTGGACACC-3’ and reverse 5’-GGTGTCCACCCAGGTAATAACTTCGTCCCTGCATAGGCT-3’ (Obtained from Eurofins Genomics).

Cells were transfected simultaneously with either non-targeting or β5 siRNA together with either EGFP alone (pEGFP-N1 empty vector; gift of Pat Caswell, University of Manchester) or WT β5-GFP using Lipofectamine 2000 (Thermofisher) according to the manufacturer’s instructions. Six hours after transfection cell cycle synchronisation was initiated by the addition of 2mM thymidine (Sigma). After 18 h the cells were released by replating in fresh medium, and a second dose of thymidine added 8 h later. The medium was replaced the next morning and imaging started approximately 5 h after the second release.

Images were acquired on an AS MDW live cell imaging system (Leica) equipped with a Cascade II EM CCD camera *(Photometrics)* and a *20x/ 0.50 Plan Fluotar air* objective. Images were collected every 10 min using Image Pro 6.3 software (Media Cybernetics Ltd) and processed using ImageJ.

#### Mitotic alignment analysis

Prior to analysis of mitotic alignment, image files were computationally blinded by randomised file name encoding. Thereafter, Fiji software was used to measure the angular difference between the long axis of the mother cell prior to cell division, and the axis of cytokinesis. All observed cell division events were recorded. Where multiple attempts at cytokinesis were observed, the orientation of the first attempt was used for angular measurement. Data were summarised using R software.

### Adhesion Assay

Cell adhesion assays were performed as described previously (Yebra et al., 1996). Briefly, non-tissue culture-treated, polystyrene 48-well cluster plates (Corning Costar Corporation) were coated with 10 μg/ml vitronectin as detailed above, and blocked with 1% heat-denatured BSA. 5 χ 10^4^ CS1-wt (negative control from non-specific attachment) or CS1-β5cells were seeded per well and allowed to attach at 37 °C for 30 min under incubation conditions. Cells were treated during attachment as indicated with combinations of cytochalasin D (20 μM) and either cyclic RAD (non-inhibitory control) or cyclic RGD (competitive inhibitor of integrin β5-vitronectin interaction) peptides (20 μg/ml), the latter as described previously (Strömblad et al., 2002). After attachment, non-adherent cells were removed by repeated washing. Remaining cells were then labeled with DAPI and imaged with a Nikon A1R confocal with 10x air objective to enable automated cell counting via NIS elements software.

## Acknowledgements

We thank David Critchley, University of Leicester, UK, for providing Talin1 null ES cells. This work was supported by grants to SS from the EU-FP7-Systems Microscopy Network of Excellence (HEALTH-F4-2010-258068), EU-H2020-Multimot (EU Grant agreement #634107), the Strategic Research Foundation of Sweden (SB16-0046), the Swedish Research Council (521-2012-3180), and the Swedish Cancer Society, and by a grant to MJH from Cancer Research UK (grant C13329/A21671). Imaging was performed at the Live cell-imaging facility and Nikon center of excellence at the Department of Biosciences and Nutrition, Karolinska Institutet, supported by grants from the Knut and Alice Wallenberg Foundation, the Swedish Research Council, the Centre for Innovative Medicine and the Jonasson donation to the School of Technology and Health, Royal Institute of Technology, Sweden. The funders had no role in study design, data collection and analysis, decision to publish, or preparation of the manuscript.

## Author contribution statement

JL’ MCJ, JAA, ML, MJH and SS conceived the project and devised experiments. JL performed live cell imaging and related image analyses. JL, MCJ and JAA performed fixed cell imaging; JL performed fixed image data analyses. XG performed the siRNA screening for PIP regulators; JL performed related image analyses. MCJ and JAA undertook experiments relating to integrin β5 RNAi and mass spectrometry analyses. AO performed STORM imaging and related image analyses. HO and SG performed immunoblotting. JL performed statistical analyses and data visualisation. JL, MCJ, JAA, XG, ML, MJH and SS contributed to writing of the manuscript.

---

**Supplementary Figure 1.**
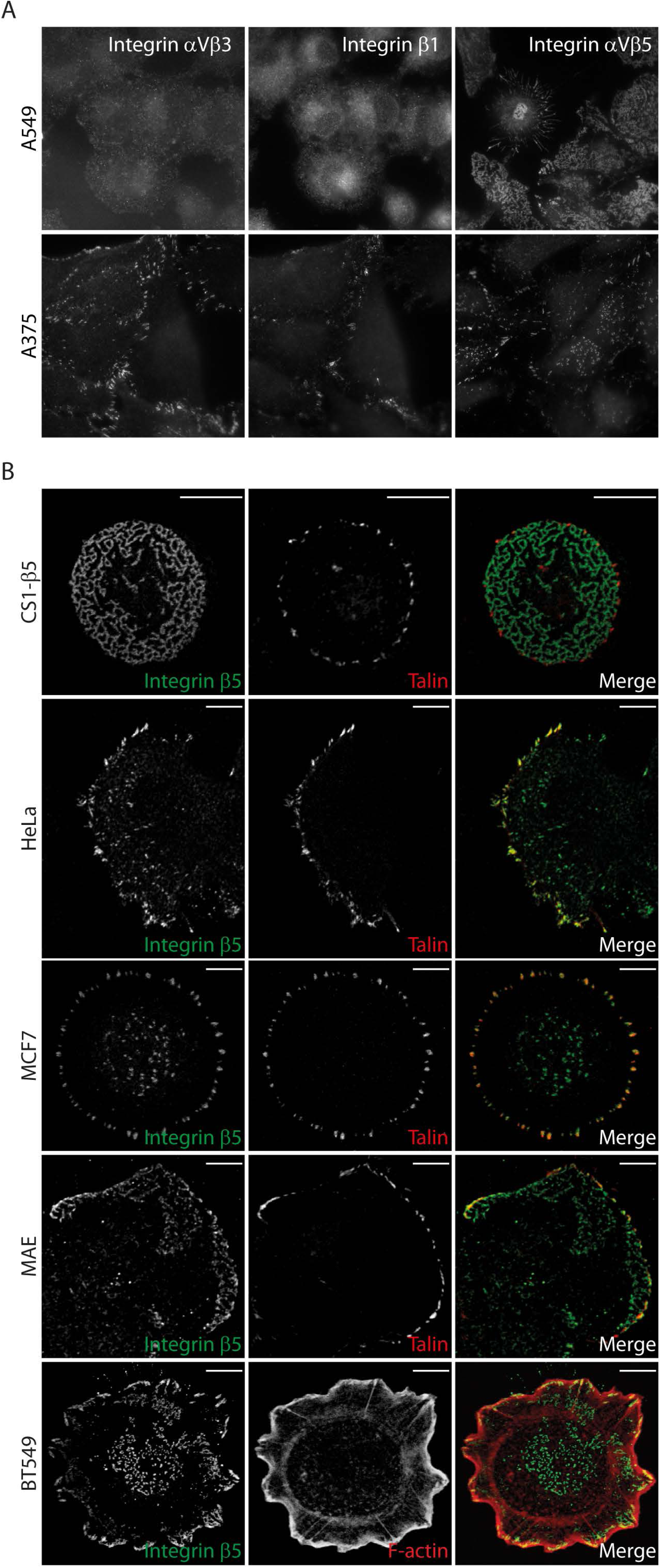
β5-positive, talin- or F-actin-negative structures found in multiple cell lines. (A) A549 lung cancer cells and A375 melanoma cells plated on glass coverslips for 72 h. Confocal images of immuno-fluorescently labeled integrins αVβ3 (LM609), β1 (9EG7) and αVβ5 (15F11). (B) Human-, mouse- and hamster-derived cell lines were found to contain central β5-positive, talin- and F-actin-negative structures equivalent to those first observed in U2OS cells. Confocal images of cells replated for 3 h on 10 μg/ml vitronectin and immuno-labeled for integrin β5 and either talin or F-actin, as indicated. Scale bars: 10 μm.

**Supplementary Figure 2.**
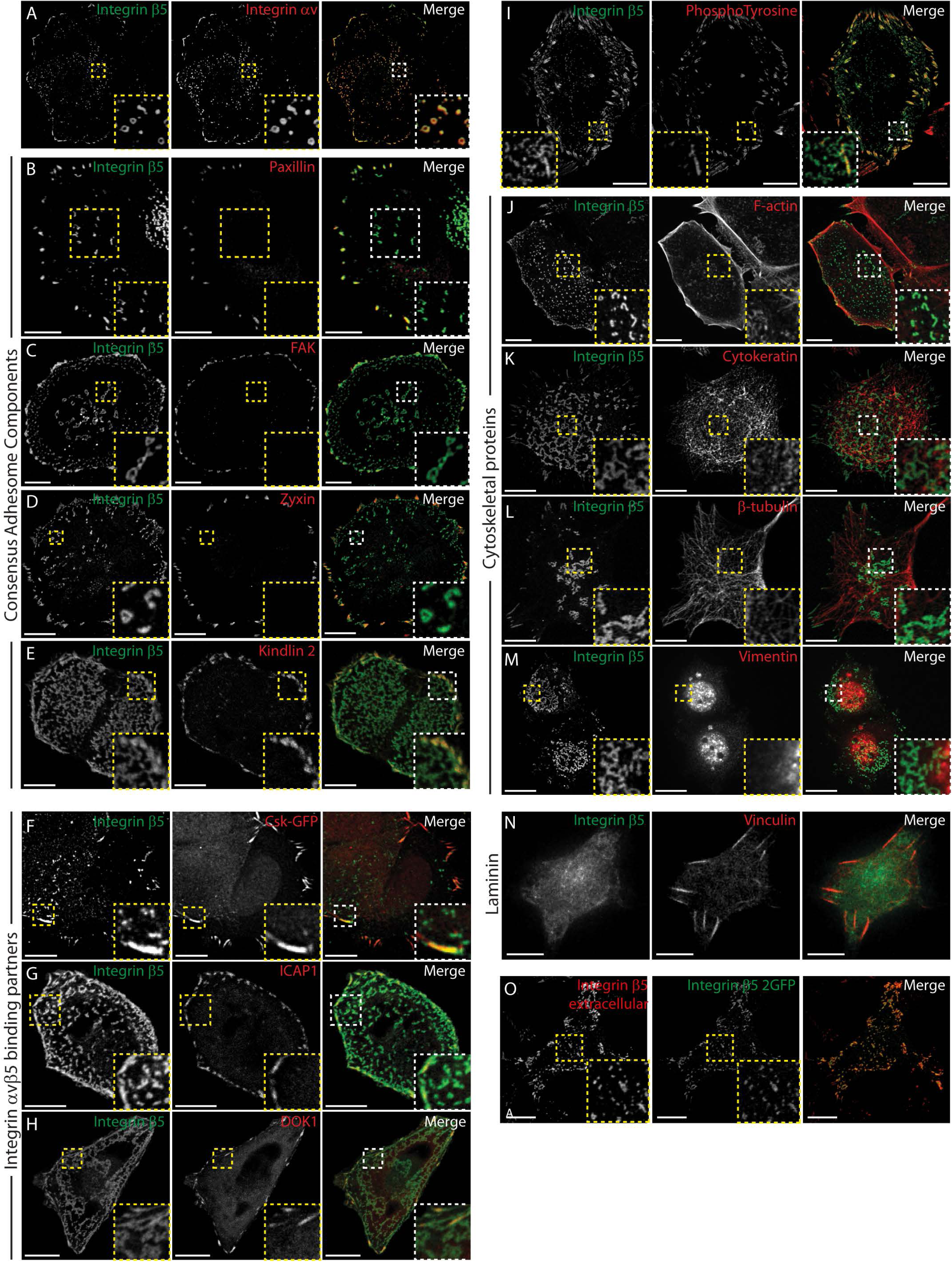
β5-positive reticular adhesions lack consensus adhesome components. (A-M) Confocal images of U2OS cells plated for 3 h on vitronectin and immuno-labeled against integrin β5 and: the alpha V (aV) subunit of the αVβ5 heterodimer (A); consensus adhesome components [paxilin (B), FAK (C), zyxin (D), kindlin 2 (E)]; integrin β 5-binding partners [CSK (F), ICAP1 (G), DOK1 (H)]; phospho-tyrosine (I); and cytoskeletal proteins [F-actin (J), cytokeratin (K), beta (β)-tubulin (L), vimentin (M)]. (N) Confocal image of U2OS cells plated for 3 h on laminin (ECM ligand not bound by αVβ5) and immuno-labeled for integrin β5 and vinculin. (O) Confocal images of an unpermeabilised U2OS cell expressing β5-2GFP, immuno-labeled for β5. Scale bars: 10 μm. Boxed areas shown at higher magnification in lower right corners.

**Supplementary Figure 3.**
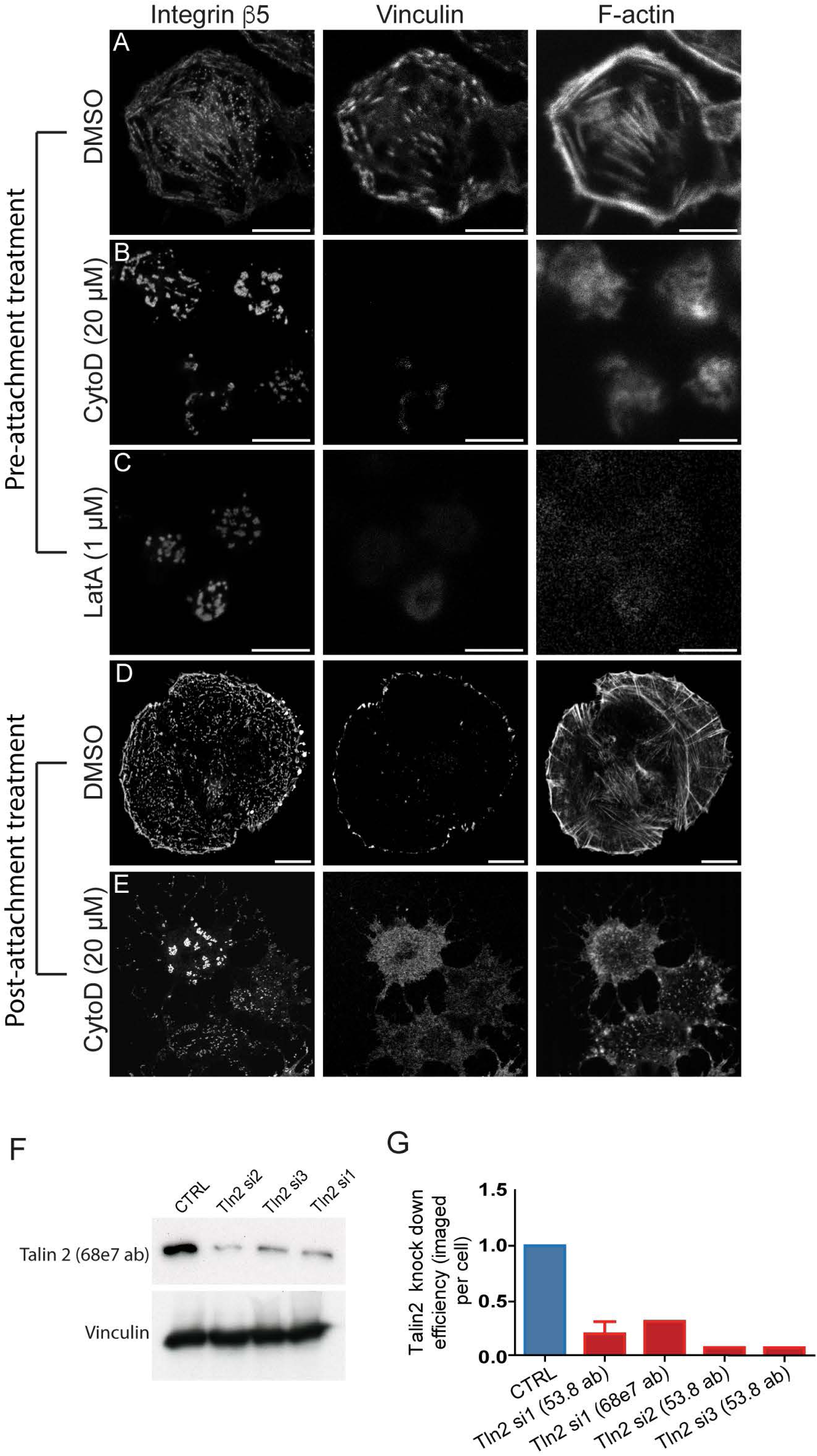
Validation of reticular adhesion independence from F-actin and talin. (A-C) Confocal images of U2OS cells attached to vitronectin (VN)-coated surfaces and labeled for integrin β5, vinculin and F-actin, following 30 min in suspension and 3 h replating in presence of DMSO (A), 20 μM cytochalasin D (CytoD) (B), or 1 μM latrunculin A (LatA) (C). Representative confocal images of U2OS cells attached to vitronectin (VN)-coated surfaces and labeled for integrin β5, vinculin and F-actin, following 3 h replating, with final 1 h in presence of DMSO (D) or 20 μM CytoD (E). (F) Representative immunoblot of talin 2 (68e7 antibody) and vinculin following siRNA treatment with control (CTRL) oligonucleotides or one of three alternative talin 2-targeting oligonucleotides. (G) Single cell imaging-based quantification of residual talin fluorescence relative to control siRNA, using the same three oligonucleotides and alternative talin 2 antibodies (53.8, 68e7). Tln2 si1 in combination with talin 2 antibody 53.8 replicated 4 times; error bars indicate standard deviation.

**Supplementary Figure 4.**
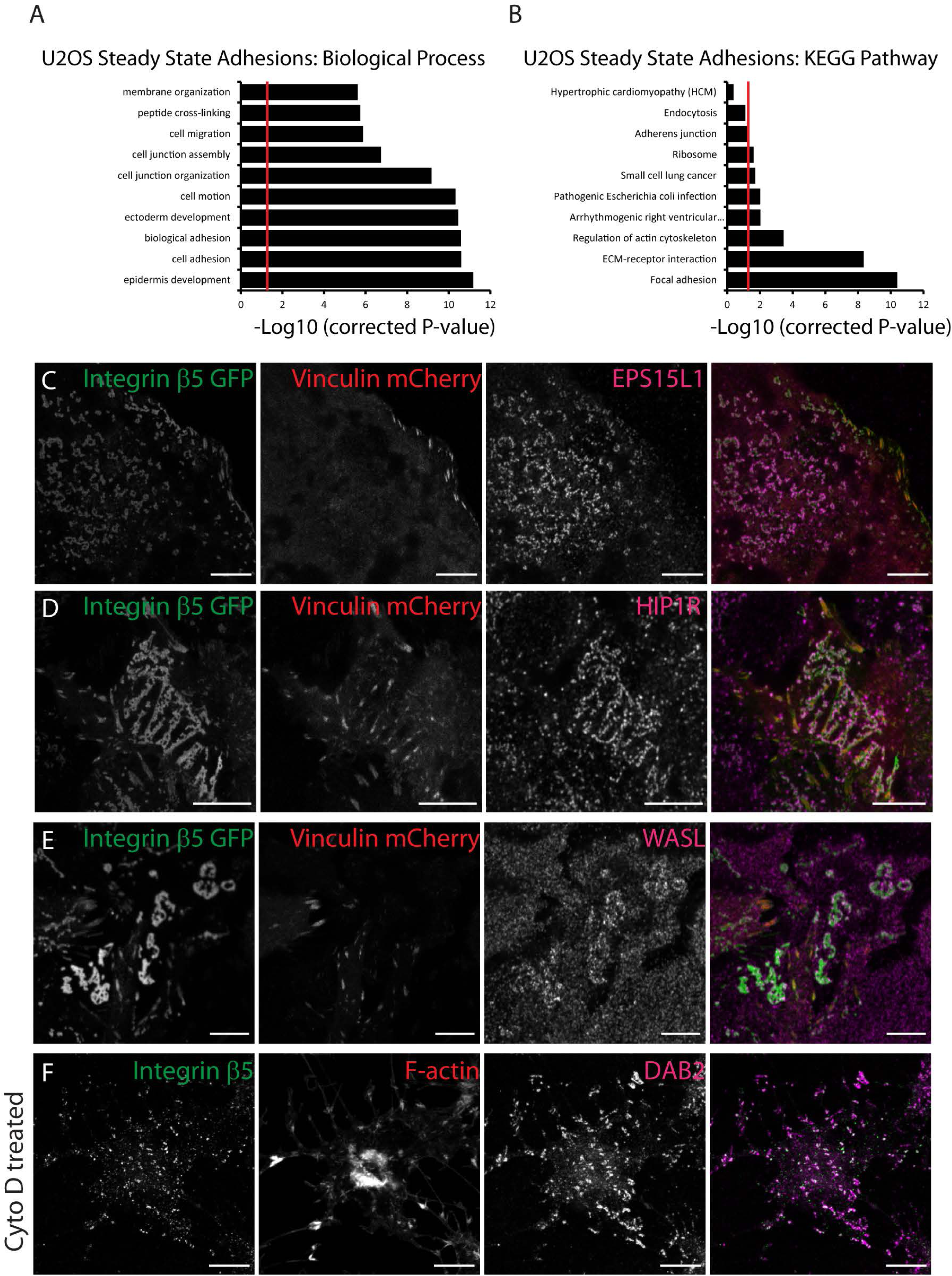
Validation of reticular adhesion proteins identified by mass spectrometry. (A,B) Gene-ontology analysis of U2OS steady state adhesion proteins (DMSO treated; focal and reticular adhesions) showing terms from Biological Process (A) and KEGG pathway analysis (B) significantly enriched over whole cell proteome. (C-E) Confocal images of U2OS β5 V cells attached to vitronectin (VN)-coated surfaces and immuno-labeled against EPS15L1 (C), HIP1R (D), or WASL (E). (F) Confocal images of U2OS cells cultured on glass coverslips for 72 h then treated with 20 μM CytoD for 2 h and immuno-labeled against integrin β5 and DAB2 along with staining of F-actin.

**Supplementary Figure 5.**
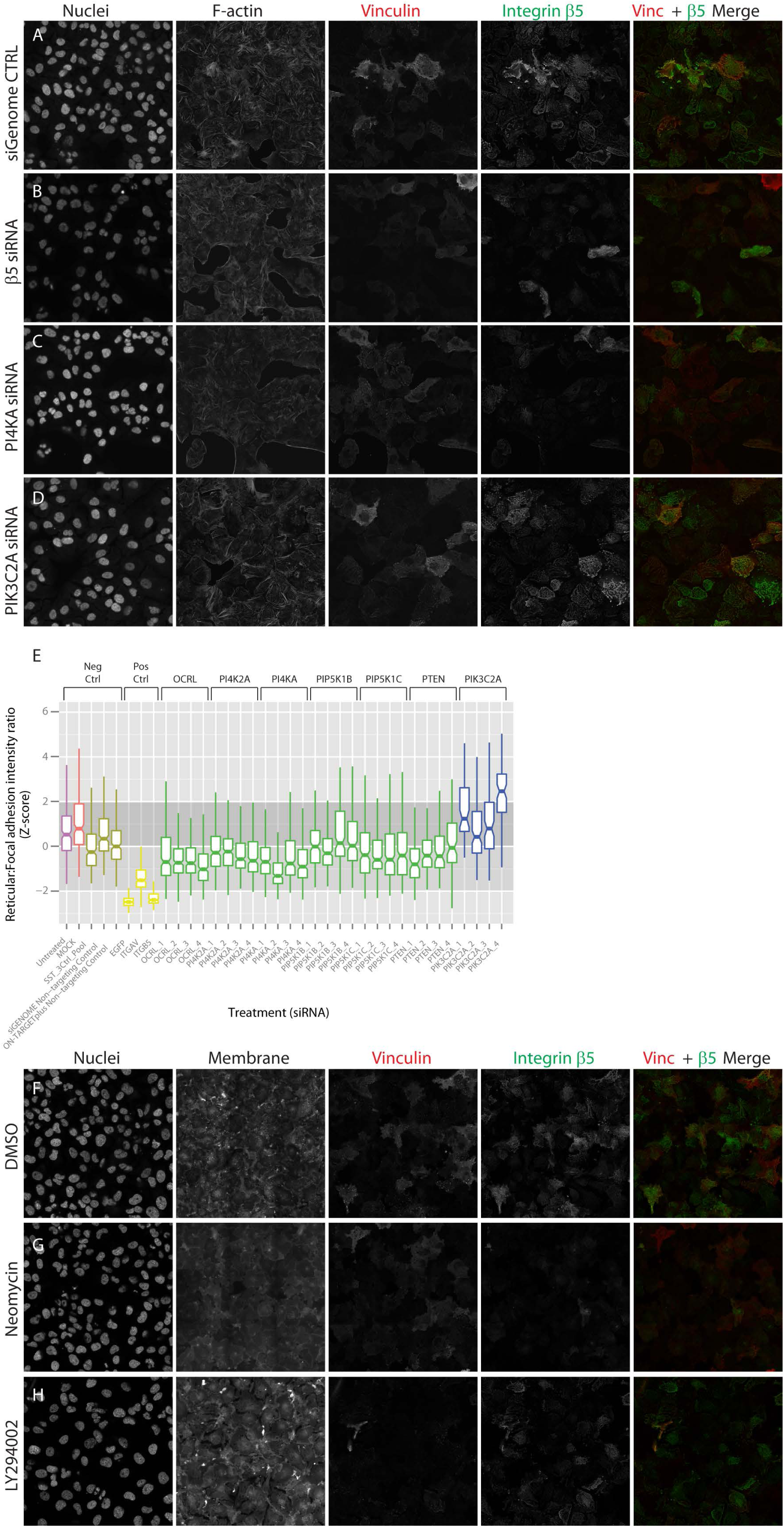
Validation of RNAi screen to identify role of PIP signalling in reticular adhesion formation. (A-D) Representative images of U2OS β5V cells stained with DAPI and phalloidin, and showing integrin β5-2GFP and vinculin mCherry fluorescence following treatment with control siRNA (A) or siRNAs targeting β5 (B), PI4KA (C) or PIK3C2A (D). (E) Quantification of reticular to focal adhesion integrin β5 ratios, per cell, for 4 independent siRNAs targeting controls or PIP regulators. (F-H) Representative images of U2OS cells stained with DAPI and phalloidin and showing integrin β5-2GFP and vinculin mCherry fluorescence following treatment with DMSO (F), 10 mM Neomycin (G) or 25 μM LY294002 (H).

**Supplementary Figure 6.**
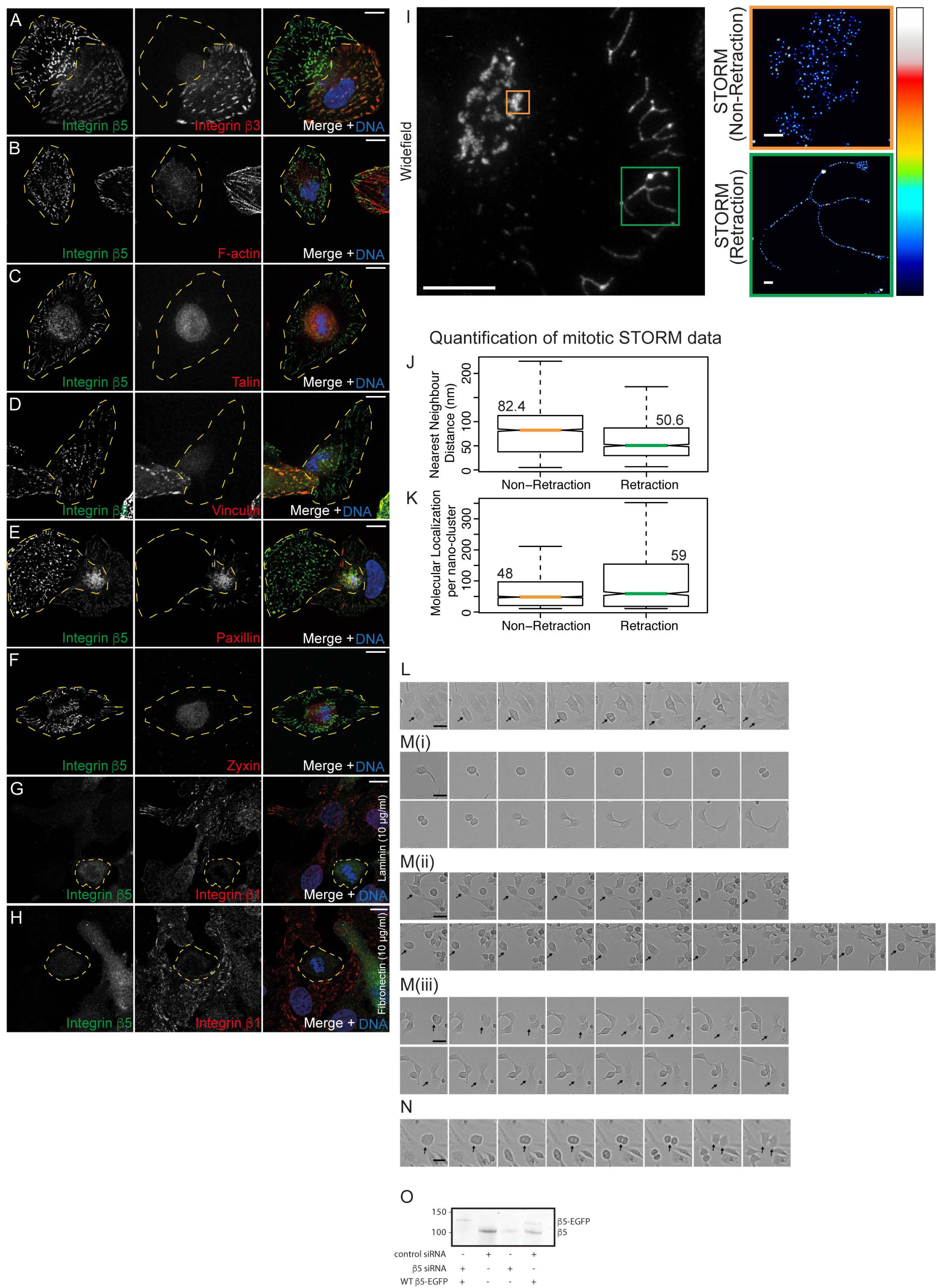
Integrin β5 reticular adhesions persist during mitosis and knockdown of β5 abrogates cell division. (A-F) Confocal images of mitotic U2OS cells (yellow dashed outlines) plated on VN and labeled for integrin β5 and: integrin β3 subunit (A); F-actin (B) and consensus adhesome components [talin (C), Vinculin (D), Paxillin (E), zyxin (F)]. (G-H) Confocal images of mitotic U2OS cells plated on 10μg/ml laminin (G) or fibronectin (H) immuno-labeled against integrin β5 and integrin β1 subunits. (I) Integrin β5 in a representative U2OS mitotic cell plated on VN and imaged via conventional (diffraction-limited) confocal microscopy. Representative central (Non-retraction) and peripheral (Retraction) reticular adhesions cropped from matched conventional and stochastic optical reconstruction microscopy (STORM, ‘royal’ look-up table scales from minimum (black) to maximum (white) intensity values as indicated in legend) images (right). (J,K) Quantification of integrin β5 nanocluster nearest neighbor distances (J) and molecular localization counts per nanocluster (K) based on STORM data. In total, 95 retraction and 83 non-retraction mitotic reticular adhesions were quantified, including > 4000 nanoclusters. Boxplot notches approximate 95% CI. Scale bars: (A-H) 10 μm (I) 2 μm (500 nm in cropped images). (L) Hela cells with control siRNA showing normal cell division (see **Supplementary Movie 10**). (M) Three example microscopy time series showing different effects of depleting β5 on cell division: i) cell remains round for an extended time before eventually dividing, often with incomplete cytokinesis (see **Supplementary Movie 11**. ii) cell repeatedly rounds up and respreads without dividing (see **Supplementary Movie 12**. iii) cell appears to divide but before cytokinesis merges back into a single bi-nucleate cell (see **Supplementary Movie 13**. (N) Hela cell treated with β5 siRNA and rescued with β5-EGFP (see **Supplementary Movie 14**). (L-N) Images were taken at 10 min intervals, scale bars = 50 μm. Arrows point to mitotic cells of interest. (O) Immunoblot of endogenous and β5-EGFP expression in control and knockdown cells.

**Supplementary Table 1.**
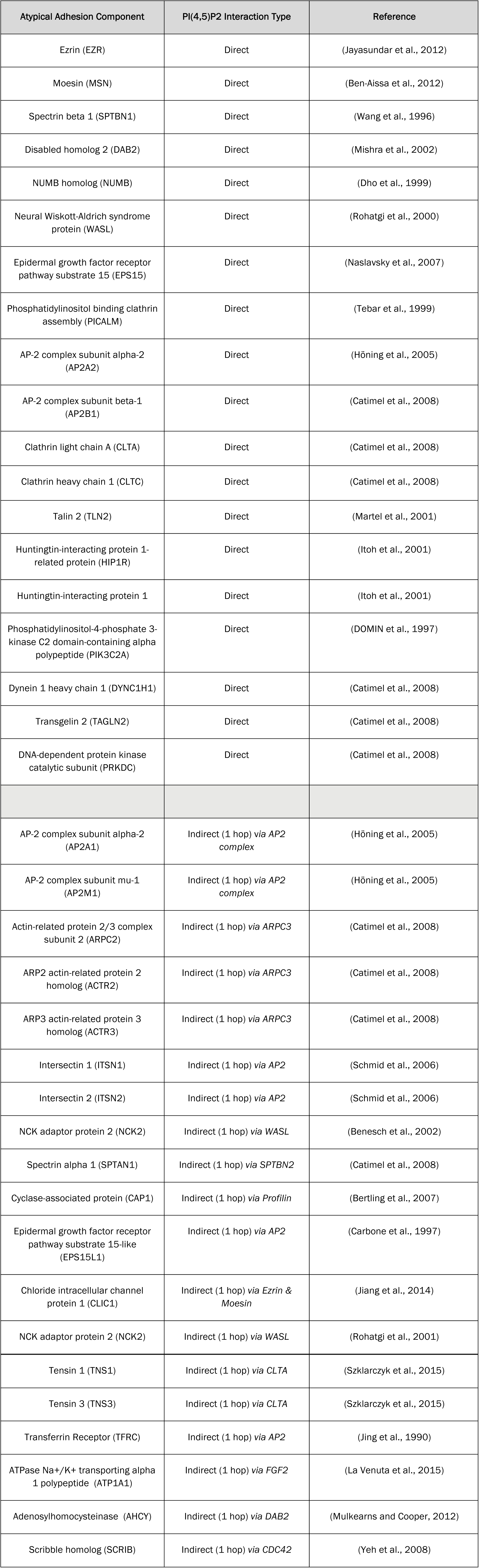
***Direct and indirect PI45P2 interactors*** This table provides references to evidence of direct or indirect interactions between reticular adhesome components and PI(4,5)P2. Indirect interactors are limited in this case to ‘1 hop’ interactions, meaning interaction through a single binding partner intermediary.

**Supplementary Table 2.**
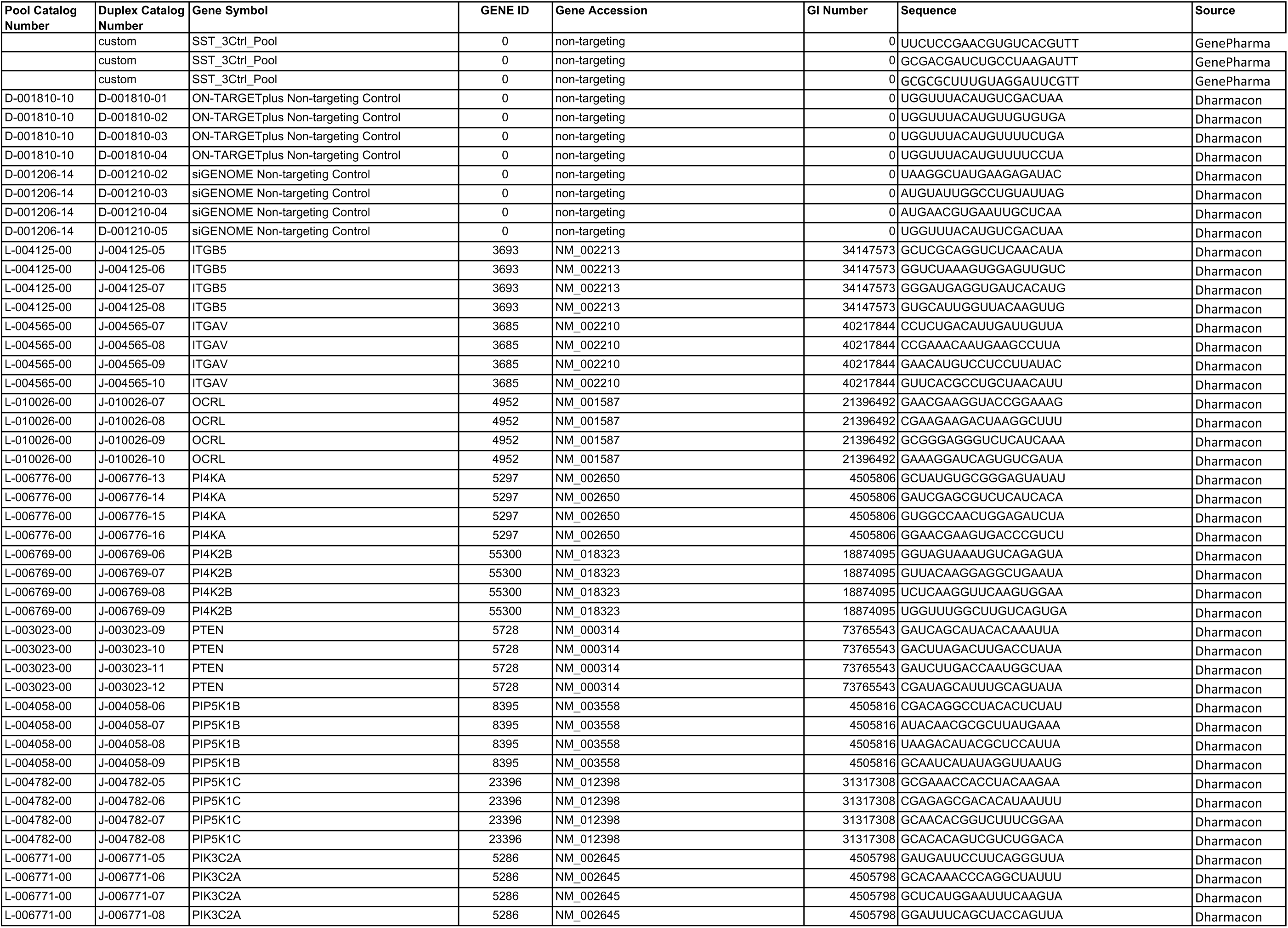
***PIP regulator siRNA screen oligonucleotide sequences*** This table provides sequences for each of the siRNA oligonucleotide sequences used in the PIP regulator screen.

***Supplementary Movie 1. Reticular adhesion formation, maturation and turnover***

Confocal live cell imaging of U2OS β5V cells at 10 min intervals for 12 h, as depicted in Figure 2E-G. Vinculin mCherry (left, red in merge), integrin β5 2GFP (centre, green in merge) and merge (right) channels demarcate reticular (vinculin mCherry-negative; typically more centrally located) and classical focal adhesions (vinculin mCherry-positive; typically peripherally located). Note that reticular adhesions form, mature and disassemble without ever showing vinculin mCherry concentration.

***Supplementary Movie 2. Reticular adhesions do not recruit vinculin during their lifetime***

Confocal live cell imaging of U2OS β5V cells at 10 min intervals for 12 h, cropped from Supplementary Movie 1 (as indicated in Figure 2E-G), as depicted in Figure 2H. Vinculin mCherry (left, red in merge), integrin β5 2GFP (centre, green in merge) and merge (right) channels. Integrin β5 2GFP demarcates the formation, maturation and disassembly of a single reticular adhesion. Note that this complete life cycle occurs without the recruitment of any detectable concentration of vinculin mCherry.

***Supplementary Movie 3. Comparison of reticular and focal adhesion dynamics***

Spinning-disc confocal live cell imaging of U2OS β5V cells at 10 min intervals for 12.5 h, as depicted in Figure 2I-M. (Upper panels) Vinculin mCherry (left, red in merge), integrin β5 2GFP (centre, green in merge) and merge (right) channels. (Low panels) Focal adhesion tracking demarcated by vinculin mCherry (left, tracks rainbow colour-coded indicating recent (blue-to-green) and old (organge-to-red) movements), focal adhesion and reticular adhesion tracking demarcated by integrin β5 2GFP (centre, tracks rainbow colour-coded indicating recent (blue-to-green) and old (organge-to-red) movements) and merged tracking results (right) channels. Note that the subset of reticular adhesion dynamics was specifically extracted for quantitative analyses by parsing tracking results with quantification of vinculin mCherry content (intensity) in integrin β5 2GFP-defined adhesions, using the criteria described in methods.

***Supplementary Movie 4. FRAP comparison of integrin β5 2GFP***

Confocal-based fluorescence recovery after photobleaching (FRAP) analysis of U2OS β5V cells to compare integrin β5 2GFP turnover rates in reticular versus focal adhesions, as depicted in Figure 2O-T and described in methods. Vinculin mCherry (left, red in merge), integrin β5 2GFP (centre, green in merge) and merge (right).

***Supplementary Movie 5. Reticular adhesion formation during cell-ECM attachment in the presence and absence of F-actin***

Confocal images of integrin β5-2GFP and vinculin mCherry in live U2OS β5V cells over 3.6 h post attachment to vitronectin-coated glass, as detailed in Figure 3. Cells were pre-treated in suspension (30 min) and during spreading with either DMSO (upper panels) or 20 μM of F-actin polymerization inhibitor cytochalasin D (CytoD; lower panels). Integrin β5 2GFP (left, green in merge), vinculin mCherry (centre, red in merge) and merge (right) channels.

***Supplementary Movie 6. Reticular adhesions persist throughout cell division when classical focal adhesions disassemble***

Spinning-disc confocal live cell imaging of U2OS β5V cells at 10 min intervals for 16.5 h, as depicted in Figure 6C-I. (Upper panels) Membrane dye (left, magenta in merge), vinculin mCherry (centre left, red in merge), integrin β5 2GFP (centre right, green in merge) and merge (right) channels. (Lower panels) Intensity color-coded (‘fire’ look-up table scales from low intensity to high via black, red, yellow, white) vinculin mCherry signal (left), and segmented and tracked vinculin-labeled focal adhesions (centre left; red outlines) and cell outline (blue outline demarcated by membrane dye). Not loss of focal adhesions during mitosis. Intensity color-coded (‘fire look-up table) integrin β5 2GFP (centre right) and segmented and tracked reticular and focal adhesions (right; red outlines) as well as cell outline (blue outline).

***Supplementary Movie 7. Mitotic retraction fibres attach to ECM at reticular adhesion sites***

3-dimensional confocal reconstruction of a U2OS β5V cell mid-mitosis, as depicted in Figure 6J-L. Membrane labelling (red) demarcates the cell body and retraction fibres angling down from the cell cortex to the ECM interface. Retraction fibres terminate in integrin β5 2GFP-decorated reticular adhesions (green). DNA is revealed in compacted form (white) via a progressive cut-through of the reconstructed view from the dorsal surface.

***Supplementary Movie 8. Post-mitotic daughter cells re-spread via retraction fibres tethered to the ECM by reticular adhesions, thereby recovering the pre-mitotic footprint***

4-dimensional confocal reconstruction of a U2OS β5V cell imaged at 10 min intervals during the respreading of daughter cells post-mitosis. Membrane labelling (red) demarcates the cell body and retraction fibres angling down from the cell cortex to the ECM interface. Retraction fibres terminate in integrin β5 2GFP-decorated reticular adhesions (green). DNA (blue) revealed via a cut-through of the reconstructed view.

***Supplementary Movie 9. Individual centrally located reticular adhesion complexed persist throughout mitosis***

Spinning-disc confocal live cell imaging of U2OS β5V cells at 10 min intervals for 13h 50 min. Integrin β5 2GFP (left, green in merge), vinculin mCherry (centre, red in merge) and merge (right) channels. Note the persistence of integrin β5 2GFP-decorated reticular adhesions throughout mitosis when classical focal adhesions labeled by vinculin mCherry disassemble. In particular, note that centrally located reticular adhesions are morphologically stable throughout mitosis, indicating that the same individual reticular adhesions persist throughout mitosis, not only the reticular adhesion population in general. This supports a role for reticular adhesion in retention of spatial memory from one cell generation the next.

***Supplementary Movie 10. Cell division in Hela cells after synchronisation***

Hela cells, transfected with control siRNA, were synchronised by double thymidine block and imaged 8h after second release. Images were taken every 10min for a further 320min. Movie shows normal cell division taking approximately 70min to complete.

***Supplementary Movies 11-13. Defects in cell division after integrin β5 knockdown***

Hela cells transfected with integrin β5 siRNA were synchronised and imaged as above. Movies show different mitotic defects that resulted. **Movie 11** cell remained rounded for an extended period of time often with incomplete cytokinesis. **Movie 12** cell repeatedly rounds up and respreads without dividing. **Movie 13** cell appeared to divide but before cytokinesis merged to form a bi-nucleate cell.

***Supplementary Movie 14. Rescue of cell division defects by wild type integrin β5***

Hela cells transfected with integrin β5 siRNA together with WT β5-EGFP were synchronised and imaged as above. Movie shows rescue of mitotic defects.

